# Learning about threat from friends and strangers is equally effective: an fMRI study on observational fear conditioning

**DOI:** 10.1101/2021.09.20.461036

**Authors:** Anna M. Kaźmierowska, Michał Szczepanik, Marek Wypych, Dawid Droździel, Artur Marchewka, Jarosław M. Michałowski, Andreas Olsson, Ewelina Knapska

**Affiliations:** Laboratory of Brain Imaging, Nencki Institute of Experimental Biology of Polish Academy of Sciences, 3 Pasteur Str., 02-093 Warsaw, Poland; Laboratory of Emotions Neurobiology, BRAINCITY - Centre of Excellence for Neural Plasticity and Brain Disorders, Nencki Institute of Experimental Biology of Polish Academy of Sciences, 3 Pasteur Str., 02-093 Warsaw, Poland; Laboratory of Affective Neuroscience, SWPS University of Social Sciences and Humanities, 10 Kutrzeby Str., 61-719 Poznań, Poland; Department of Clinical Neuroscience, Division of Psychology, Karolinska Institutet, Stockholm, Sweden

**Keywords:** fear contagion, observational fear learning, familiarity, social learning, ecological validity

## Abstract

Humans often benefit from social cues when learning about the world. For instance, learning about threats from others can save the individual from dangerous first-hand experiences. Familiarity is believed to increase the effectiveness of social learning, but it is not clear whether it plays a role in learning about threats. Using functional magnetic resonance imaging, we undertook a naturalistic approach and investigated whether there was a difference between observational fear learning from friends and strangers. Participants (observers) witnessed either their friends or strangers (demonstrators) receiving aversive (shock) stimuli paired with colored squares (observational learning stage). Subsequently, participants watched the same squares, but without receiving any shocks (direct-expression stage). We observed a similar pattern of brain activity in both groups of observers. Regions related to threat responses (amygdala, anterior insula, anterior cingulate cortex) and social perception (fusiform gyrus, posterior superior temporal sulcus) were activated during the observational phase, reflecting the fear contagion process. The anterior insula and anterior cingulate cortex were also activated during the subsequent stage, indicating the expression of learned threat. Because there were no differences between participants observing friends and strangers, we argue that social threat learning is independent of the level of familiarity with the demonstrator.

**Highlights:**

- We compared observational learning of fear from friends and strangers
- Familiarity does not enhance social learning of fear in humans
- Bayesian statistics confirm absence of differences between friends and strangers
- Observational fear learning activates social and fear networks including amygdala
- Amygdala activations are absent when learned fear is recalled

## 1. Introduction

Detecting potential threats in the environment is vital for survival. To learn about danger, social species, including humans, use direct and vicarious cues (Olsson et al., 2020). Using vicarious cues is beneficial from the evolutionary perspective as it might eliminate the risk of first-hand encounters with a threat. One of the mechanisms underlying vicarious learning about dangers is emotional contagion, an automatic, unconscious tuning into the affective state of others. Due to such emotional mimicry, the observers can ‘feel’ the threat while enjoying relative safety themselves. Through exposure to the observed danger, they can learn about the potential risks within the environment.

According to the well-established Russian doll model of empathy, the strength of the relationship between the ‘demonstrator’ and the ‘observer’ modulates emotional contagion (Preston & de Waal, 2002). The model predicts that emotional contagion is more robust among individuals in close social relationships as their perception-action coupling is more powerful, and empathy is higher. Empirical studies have confirmed that in general, familiarity improves the social transfer of emotions (Bruder et al., 2012; de Vignemont & Singer, 2006; Gonzalez-Liencres et al., 2014; Hatfield et al., 2014). Importantly, however, previous research suggests that a close social relationship does not facilitate the emotional convergence of fear (Bruder et al., 2012). The only report comparing the neural correlates of a threat (an electric shock) to self, a friend, and a stranger gave no conclusive results (Beckes et al., 2013). Besides no significant difference found in a direct friend-stranger comparison, the within-subject design employed in this study did not allow for inferring about the individual effect of different experimental conditions. Thus, the impact of familiarity on fear contagion and observational fear learning remains unclear.

To address this open question, we used a modified observational fear learning protocol (Haaker, Golkar, et al., 2017) improved in terms of ecological validity (Szczepanik, Kaźmierowska, et al., 2020). We directly tested how the demonstrator’s familiarity influenced both psychophysiological (skin conductance response, SCR) and brain (functional magnetic resonance imaging, fMRI) correlates of observational fear learning in the observer. We measured SCR to include a modality used in previous studies on observational fear learning (Golkar et al. 2015; Golkar and Olsson 2017; Haaker et al. 2017) and validate our paradigm (Szczepanik, Kaźmierowska, et al., 2020), as well as to enhance the power of our inferences. The experimental design (Figure 1) involved two main stages. During the observational learning stage, two groups of participants (observers) witnessed their friends or strangers (demonstrators) undergoing a differential fear conditioning procedure. In this task, uncomfortable but not painful electric shocks were applied to the demonstrator’s forearm during the presentation of emotionally neutral visual stimuli. Subsequently, during the direct-expression stage, we tested the observers’ brain responses to previously watched stimuli. No electrical stimulation was applied to the observers throughout the whole experiment. Thus, the observers derived the knowledge about the aversive value of stimuli only from social learning. As we wanted our experimental situation to be as naturalistic as possible, no apparent ingroup-outgroup division was introduced (we did not emphasize the demonstrator’s affiliation). Also, unlike most previous studies that have used an actor serving as the demonstrator, real-time videos were transmitted to the observers in the friend-observation condition. In the stranger-observation condition, we used recordings from the friend-observation condition. Developing such an ecologically valid paradigm enabled investigating social learning of fear in a setting relevant for naturally occurring behaviors.

**Figure 1:**
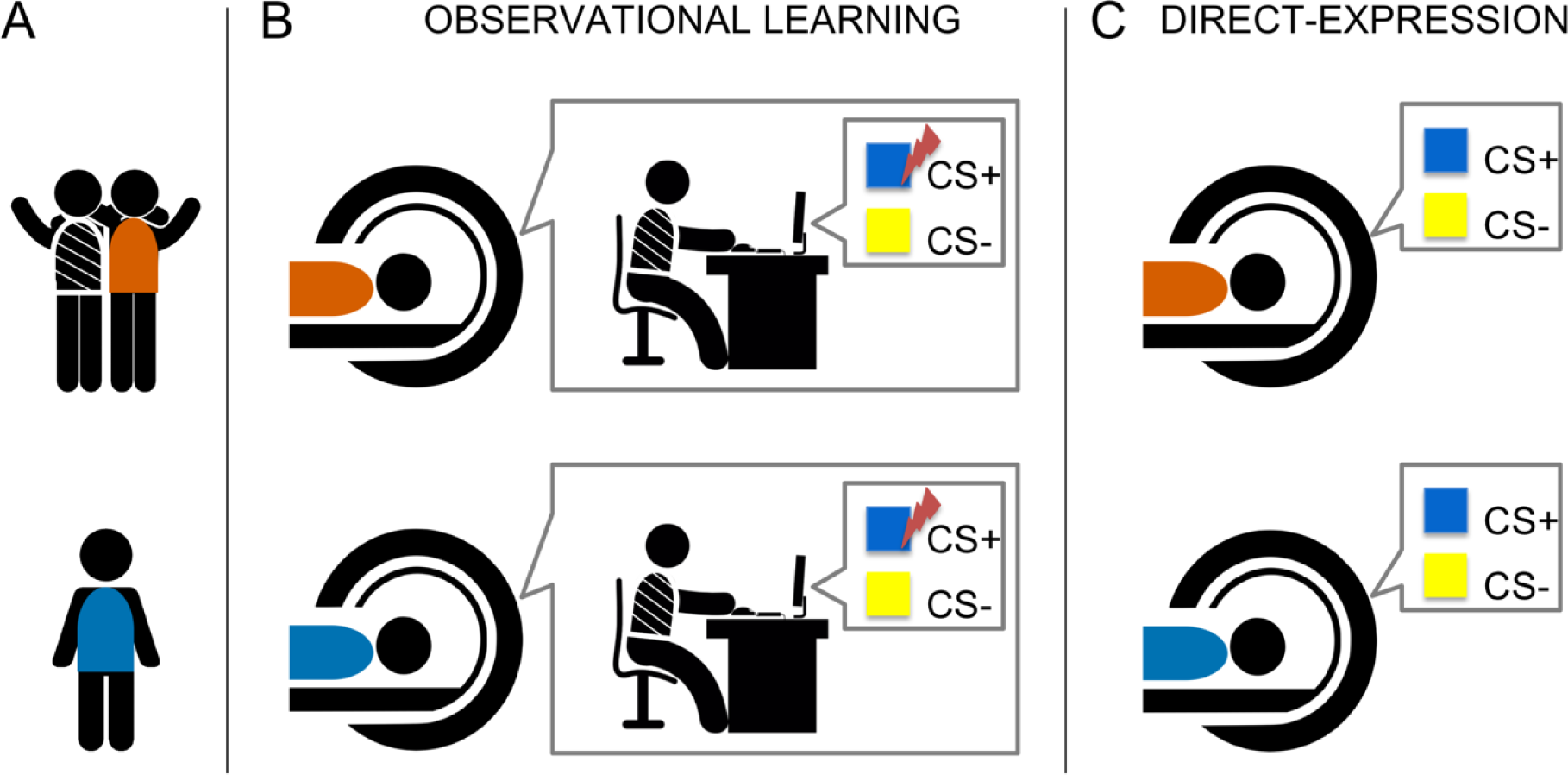
Experimental Design Note. (A) Depending on the group, observers arrived at the laboratory with their friend (upper panel, the ‘friend’ group, orange) or alone (lower panel, the ‘stranger’ group, blue). (B) During the observational learning stage, observers were lying in the fMRI scanner and watching the demonstrator (a friend or a stranger) performing the differential fear conditioning task. In the task, one stimulus (conditioned stimulus +; CS+) was sometimes paired with an uncomfortable electric shock (unconditioned stimulus, US), while the other one (conditioned stimulus -; CS-) was always safe. (C) During the direct-expression stage, observers remained in the scanner and were told that they would perform the same task as previously watched. They watched an identical set of visual stimuli, but no electrical stimulation was applied.

Based on the Russian doll model of empathy, we hypothesized that a more powerful perception-action coupling in friends sharing past experiences and/or higher level of empathy in the group of friends than strangers would lead to enhanced fear contagion. Thus, observing a friend’s fear will result in more robust activation in the fear network (including the amygdala, anterior insula, and anterior mid-cingulate cortex) than observing a stranger’s fear. We also predicted that the friend-observers would have a higher observational fear learning efficacy (indicated by more robust threat responses during the direct-expression stage) compared to the stranger-observers. We expected that the group differences in brain activations would be accompanied by the analogous differences in physiological responses (SCR).

## 2. Methods

### 2.1. Participants

48 pairs of friends (‘friend’ group) and 47 individuals (‘stranger’ group) participated in the experiment. Pairs of friends took part in the experiment together, with one participant being a demonstrator and another an observer. The demonstrators undergoing fear conditioning were live video streamed in the ‘friend’ group, and their video recordings were used in the ‘stranger’ group. Only the observers underwent fMRI. To estimate groups’ sizes that provide sufficient statistical power, we considered a similar fMRI experiment with the between-group design (n=21 and n=22 (Haaker et al., 2017)). We increased the number of recruited individuals as the rate of contingency-aware participants is lower in a real-time demonstrator-observer interaction (Szczepanik, Kaźmierowska, et al., 2020). The eligibility criteria included being heterosexual male, aged between 18 and 30 years, right-handed, and native or fluent Polish speaker. Only heterosexual participants (based on self-declaration) were recruited to restrict the relationships to non-romantic male friendship to reduce sample variability. Handedness was assessed on declarative level, both in the recruitment and fMRI safety-screening forms. We excluded students and graduates of either psychology or cognitive science, participants with neurological disorders or other medical conditions precluding MR scanning or electrical stimulation, and participants taking psychoactive drugs. Additionally, in the ‘friend’ group, the participants had to have known each other for at least three years and score at least 30 out of 60 points in the McGill Friendship Questionnaire - Respondent’s Affection (Mendelson & Aboud, 1999), see section 2.4.1.

We excluded four subjects from the ‘stranger’ group: three subjects who experienced technical problems with the video playback, and one subject who showed excessive head motion during one of the tasks (more than 25% volumes classified as motion outliers, see fMRI Data Preprocessing section). We included only the contingency-aware participants in the analyses; please refer to section 2.4.4. The final group sizes were n = 35 (‘friend’ group) and n = 34 (‘stranger’ group). In the analyzed sample, the mean age of all observers was 22.9 years (SD = 2.87), the mean length of the observer-demonstrator friendship was 7.7 years (SD = 4.22), and the mean observers’ score in the McGill Friendship Questionnaire was 52 (SD = 7.74), see Table S2 in Appendix for the detailed statistics. All participants received financial remuneration of 100 PLN (∼23 EUR) for their participation.

### 2.2. Tasks and Stimuli

#### 2.2.1. Experimental Setup

In the MRI scanner, the observer watched a video (either a streaming or a replay, both without sound) or picture stimuli on an MR-compatible monitor through a mirror box placed on the head coil. The demonstrator sat in a small room adjacent to the MR room (in the ’friend’ group). A GoPro Hero7 camera provided video transmission and recording. To ensure reliable reproduction of stimuli’ colors, we lighted the room with an LCD panel with adjustable color temperature, set the computer screen brightness to low, and adjusted the camera’s white balance. The room walls were covered with gray acoustic foam to minimize distractors.

#### 2.2.2. Stimuli

The conditioned stimuli were two squares, blue and yellow, displayed on a gray background. The squares were presented centrally and covered either a half (during the observation stage) or a quarter (during the direct-expression stage) of screen height (Figure 2). The assignment of color to CS+ and CS- was counterbalanced across participants.

**Figure 2:**
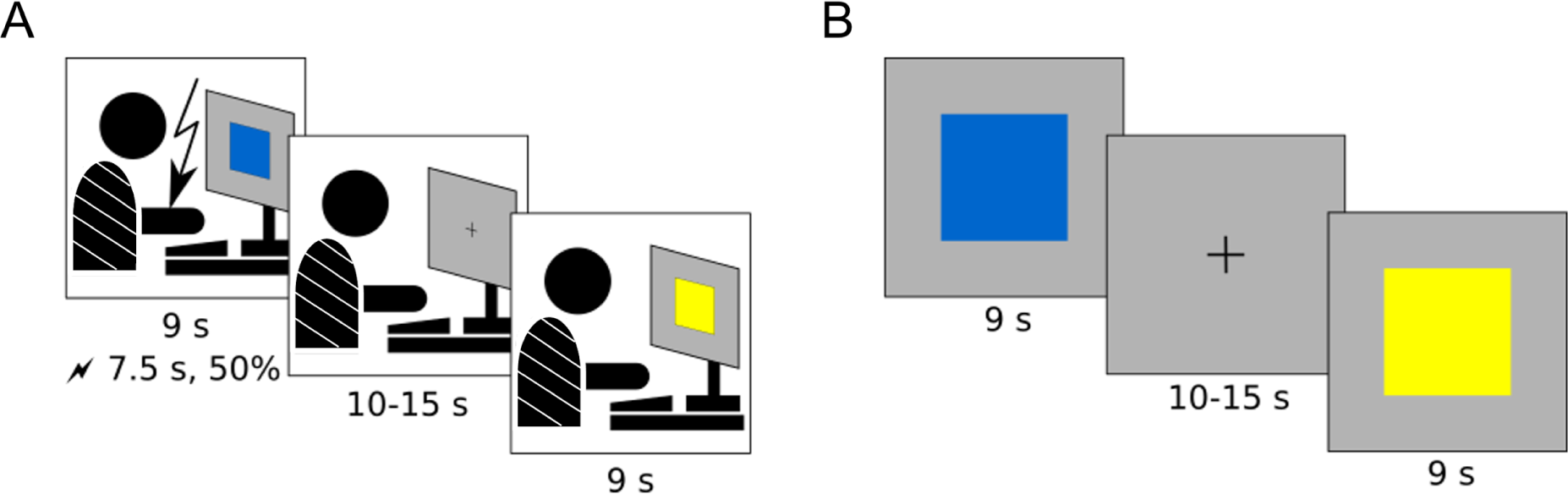
Tasks and Stimuli Overview Note. Schematic representation of the task design. (A) The observational learning stage: participants watched another person (demonstrator) who performed a differential conditioning task. The US (electrical stimulation) reinforced half of the visual CS+, CS- were never reinforced. (B) The direct-expression stage: participants, informed that they would undergo an identical task as the demonstrators, watched the same visual stimuli presentation. However, the electric stimulation was not applied.

Cutaneous electrical stimulation, applied to the ventral part of a forearm of the demonstrator, was used as the unconditioned stimulus. Stimulating electrodes were placed above the *flexor carpi radialis* muscle so that even low-intensity stimulation caused involuntary muscle flexion, visible to the observer. The stimulation consisted of five unipolar pulses of 1 ms duration applied in 200 ms intervals. The demonstrators individually adjusted shock intensity to be unpleasant but not painful (see section 2.3.3).

#### 2.2.3. Observational Learning Stage

The observer watched the demonstrator performing a differential conditioning task. The demonstrator watched 24 CS+, and 24 CS- displayed on the screen in pseudo-random order, with a given CS repeated at most twice in a row. Each CS lasted 9 seconds, half of the CS+ reinforced with the US. The US reinforced the first and the last presentation of CS+. The US started 7.5 seconds after the CS onset to allow the demonstrator’s reaction to co-terminate with the CS. The CS- was never reinforced. The intertrial intervals (ITIs) lasted randomly between 10 and 15 seconds, with a fixation symbol (+) displayed centrally on the screen (Figure 2).

#### 2.2.4. Direct-expression Stage

The observer watched 12 CS+ and 12 CS-. The stimuli and the stimuli’ order, timing, and ITI were the same as during the observational learning stage. None of the CS was reinforced.

### 2.3. Procedure

To investigate observational fear learning, we adapted the protocol of Haaker, Golkar, et al. (2017) for live observation of demonstrator-observer interaction (Szczepanik, Kaźmierowska, et al., 2020). The protocol was approved by the Ethics Committee of the Faculty of Psychology at the University of Warsaw (decision from 28 November 2017). The procedure complied with the Ethical Principles of Psychologists and Code of Conduct published by the Polish and the American Psychological Associations.

#### 2.3.1. Before the Experiment

The participants received brief information about the experimental procedure, including the possibility of receiving aversive electrical stimulation. Next, the participants gave informed consent (including optional permission for recording and recording’s reuse) and filled in safety screening forms. The roles of a demonstrator and an observer were then randomly assigned to the participants by giving two color-coded envelopes. Then the participants were isolated; the demonstrators went to a room adjacent to the MR room.

#### 2.3.2. Observer’s Preparation and Instructions

First, the observers filled in the questionnaires (see section 2.4 for details). Next, they opened the envelope with the role assignment and received instructions. The instruction included information on the two-stages procedure in the fMRI scanner. The instruction said that, firstly, the participants would watch their friends perform a task involving the presentation of colored symbols and the administration of unpleasant electrical stimulation. Then, they would perform an identical task themselves. Importantly, no information about the stimuli contingency was provided. After receiving the instruction, the observers had stimulation and skin conductance electrodes attached and went to the MR room. In the scanner, the subjects had sham leads connected to the stimulation electrodes attached to make receiving electrical stimulation believable.

#### 2.3.3. Demonstrator’s Preparation and Instructions

First, the demonstrators filled in the questionnaires (see section 2.4 for details). Next, they opened the envelope with the role assignment and received instructions. The instruction said they would perform a task involving the presentation of colored symbols and administering unpleasant, but not painful, electrical stimulation. Unlike the observers, the demonstrators received information on the stimuli contingency and reinforcement rules. We asked the demonstrators to react to the stimulation in a natural yet noticeable manner. The demonstrators watched a recording with a model reaction. After receiving the instruction, the demonstrators had stimulation and sham skin conductance electrodes attached. Afterward, the demonstrators adjusted stimulation intensity. The experimenter increased it stepwise, and the participants rated the intensity using a scale from 1 (*imperceptible*) to 8 (*painful*). The target level was 6 (*very unpleasant but not painful*).

#### 2.3.4. MRI Procedure

The MRI scanning started with the acquisition of an anatomical image. Then, the observational learning stage followed (see section 2.2.3). The experimenter adjusted the camera’s position to ensure that the observer could see the demonstrator’s face, hand, and computer screen and turned on video transmission and recording. The observer received a brief reminder: ‘in this part of the study, you will observe your friend performing a certain task’. After completion of the observational learning stage, the observer received a brief reminder: ‘in this part of the study, you will perform the same task that you just watched’, and the direct-expression stage ensued (see section 2.2.4).

#### 2.3.5. Conclusion of the Experiment

While the observer took part in the direct-expression stage, the demonstrator completed the post-experimental questionnaires (see section 2.4). After the MRI session ended, the observer also completed their set of post-experimental questionnaires, and both participants were debriefed about the study.

#### 2.3.6. ‘Stranger’ Group

In the ‘stranger’ group, only one subject (the observer) participated in the experimental session. Instead of a live stream, we used 45 recordings made in the ‘friend’ group. Otherwise, the procedure in the ‘stranger’ group was very similar to that involving a pair of friends and differed only in the necessary aspects. There was no role assignment, and instructions given to the observers referred to another person rather than a friend.

### 2.4. Behavioral Measures

#### 2.4.1. McGill Friendship Questionnaire - Respondent’s Affection

To recruit pairs of friends, we used the McGill Friendship Questionnaire - Respondent’s Affection (Mendelson & Aboud, 1999); translated by A. Kaźmierowska, P. Pączek, and A. Schudy for online screening in this study. It consists of 16 positive statements describing feelings for a friend and satisfaction with the friendship, rated along a 9-point scale ranging from -4 (*very much disagree*) to 4 (*very much agree*). One item (no. 9) was omitted from the questionnaire due to human error. A score of 30 points was a threshold for inclusion in the study.

#### 2.4.2. Empathic Sensitiveness Scale

To measure the participants’ empathy, we used the Empathic Sensitiveness Scale (original title: Skala Wrażliwości Empatycznej, SWE (Kaźmierczak et al., 2007)). It is a self-report questionnaire based on the Interpersonal Reactivity Index (Davis, 1983), but with far-reaching modifications developed in the Polish language. It contains 28 statements, which reflect three components of empathy: two emotional (empathic concern, personal distress) and one cognitive (perspective taking). The answers are given on a 5-point scale. The demonstrators and observers filled in the questionnaire at the end of the experimental procedure.

#### 2.4.3. State and Trait Anxiety Inventory

To measure participants’ anxiety, we used the State and Trait Anxiety Inventory (Spielberger et al., 1983, polish adaptation 2012). It is a self-report questionnaire consisting of two 20-item subscales, STAI-State and STAI-Trait. Participants rate statements related to how they feel either at a given moment (state) or usually (trait) using 4-point Likert scales. Each participant completed the STAI-state twice (at the beginning and the end of the experiment) and STAI-trait once (at the end of the experiment).

#### 2.4.4. Assessment of Stimulus Contingency Awareness

To assess the observers’ declarative knowledge of the CS/US contingency, we used an online questionnaire adapted from (Weidemann et al., 2016). The questionnaire referred to the observational learning stage and was divided into several parts to eliminate guessing. Firstly, it asked whether the participant could predict shocks to the demonstrator. Then, if answered positively, an open-ended question ‘how’ followed. Next, participants gave percentage ratings of shock occurrence for each stimulus. Finally, they made a forced choice of one stimulus paired with the shock. CS+, CS- and a fixation symbol were all presented as stimuli in the latter two questions. Observers completed this questionnaire after the direct-expression stage to avoid interfering with the learning process (Haaker, Golkar, et al., 2017).

#### 2.4.5. Evaluation of the Demonstrator’s Expression (the Observational US)

To control how the observers perceived the demonstrators’ behavior, we used a set of questions suggested by Haaker, Golkar, et al. (2017). At the end of the experiment, we asked the observers to rate: how much discomfort the demonstrator experienced when receiving the electrical stimulation, how expressive the demonstrator was, how natural the demonstrator’s reactions were, and how much empathy they felt for the demonstrator. Additionally, we asked the observers to rate the degree of unpleasantness attributed to the demonstrators. Finally, in the ’friend’ group, the observer scored the degree to which they identified with their friend. This question was not asked in the ‘stranger’ group due to the difficulties in understanding the concept of identification with strangers that we observed during the pilot study. All ratings used a 10-point Likert scale, ranging from 0 (*not at all*) to 9 (*very much*), except for the unpleasantness rating, which used a 5-point Likert scale ranging from 1 (*very unpleasant*) to 5 (*very pleasant*).

### 2.5. Skin Conductance Response Recordings

During fMRI sessions, we registered skin conductance responses as a secondary measure. It was recorded using BrainVision BrainAmp ExG MR amplifier with GSR MR sensor, sampled at 250 Hz.

### 2.6. fMRI Data Acquisition

The MRI data were acquired on a 3T Siemens Magnetom Trio scanner equipped with a 12-channel head coil. At the beginning of a session, a T1-weighted anatomical image was acquired using an MPRAGE sequence with 1 × 1 × 1 mm resolution and the following parameters: inversion time TI = 1100 ms, GRAPPA parallel imaging with acceleration factor PE = 2, acquisition time TA = 6 minutes and 3 seconds. After acquiring the anatomical scans, two functional imaging runs followed (observational learning and direct-expression tasks). The first run contained 362 volumes in the ‘friend’ group and 380 volumes in the ‘stranger’ group. The scanning started earlier in the ’stranger’ group, but the leading volumes were removed from the analysis. The second run of scanning contained 184 volumes. Each functional volume comprised 47 axial slices (2.3 mm thick, with 2.3 × 2.3 mm in-plane resolution and 30% distance factor) that were acquired using a T2*-sensitive gradient echo-planar imaging (EPI) sequence with the following parameters: repetition time TR = 2870 ms, echo time TE = 30 ms, flip angle FA = 90 degrees, field of view FoV = 212 mm, matrix size: 92 × 92, interleaved acquisition order, GRAPPA acceleration factor PA = 2.

### 2.7. fMRI Data Preprocessing

The fMRI data were preprocessed using fMRIPrep 1.4.0 (Esteban, Blair, et al., 2019; Esteban, Markiewicz, et al., 2019), based on Nipype 1.2.0 (Gorgolewski et al., 2011; Gorgolewski et al., 2019) and Nilearn 0.5.2 (Abraham et al., 2014). We followed the fMRIprep preprocessing with smoothing in SPM (SPM 12 v7487, Wellcome Centre for Human Neuroimaging, London, UK). At the beginning of the fMRIprep pipeline, the anatomical images were corrected for intensity non-uniformity with N4BiasFieldCorrection (Tustison et al., 2010), distributed with ANTs 2.2.0 (Avants et al., 2008), and used as an anatomical reference. The anatomical reference was then skull-stripped with antsBrainExtraction (from ANTs) and segmented using fast from FSL 5.0.9 (Zhang et al., 2001). Finally, the anatomical images were normalized to the MNI space through nonlinear registration with antsRegistration (ANTs 2.2.0). The ICBM 152 Nonlinear Asymmetrical template version 2009c was used (Fonov et al., 2009).

The functional images from the observational learning and direct-expression stages were preprocessed in the following manner. First, a reference volume was generated using a custom methodology of fMRIPrep. This reference was co-registered to the anatomical reference using flirt from FSL 5.0.9 (Jenkinson & Smith, 2001) with the boundary-based registration cost-function (Greve & Fischl, 2009). Head-motion parameters with respect to the functional reference volume (transformation matrices and six corresponding rotation and translation parameters) were estimated before any spatiotemporal filtering using mcflirt from FSL 5.0.9 (Jenkinson et al., 2002). The functional scanning runs were slice-time corrected using 3dTshift from AFNI 20160207 (Cox & Hyde, 1997). Next, the functional images were resampled into the MNI space. All resamplings were performed in a single interpolation step (composing head-motion transform matrices and co-registrations to anatomical and output spaces) using antsApplyTransforms (ANTs). Finally, we smoothed the functional images with a 6 mm FWHM 3D Gaussian kernel using spm_smooth (SPM 12). Framewise displacement (FD) and the derivative of root mean square variance over voxels (DVARS) were calculated by fMRIPrep for each functional scan, both using their implementations in Nipype and following the definitions by (Power et al., 2014). Frames that exceeded a threshold of 0.5 mm FD or 1.5 standardized DVARS were annotated as motion outliers. For more details on the fMRIprep pipeline, see fMRIPrep’s documentation at https://fmriprep.org/en/latest/workflows.html.

#### Physiological Data Analysis

To analyze skin conductance data we used PsPM 4.3.0 software (https://bachlab.github.io/PsPM/) running under MATLAB 2018b (MathWorks, Natick, MA, USA). We used a non-linear model for event-related SCR. In this analysis, three parameters of sudomotor nerve response function (its amplitude, latency and dispersion) are estimated to best match the observed skin conductance data. This model is suitable for anticipatory responses in fear conditioning and assumes that the response onset following CS presentation is not precisely known (Bach et al., 2010; Staib et al., 2015). Before the analysis, we visually inspected the signals for artifacts and manually marked missing epochs. We excluded subjects with missing data or excessive presence of artifacts precluding further analysis. As a result, we analyzed SCR data for 27 subjects per group. We used the default settings for preprocessing: signals were filtered using bi-directional 1st order Butterworth filters, with 5 Hz low-pass and 0.0159 Hz high-pass cut-off frequencies, and resampled to 10 Hz. We performed no response normalization. We accelerated the processing of multiple subjects using GNU Parallel v.20161222 (Tange, 2011).

### 2.8. fMRI Data Analysis

#### 2.8.1. Activation Analysis

To analyze data we used a mass univariate approach based on a general linear model. We used SPM 12 software (SPM 12 v7487, Wellcome Centre for Human Neuroimaging, London, UK) running under MATLAB 2020a (MathWorks, Natick, MA, USA). First-level models contained four types of events in the observational learning stage: CS+, CS-, US, and no US (i.e., lack of US during 50% of CS+). The observational CS were modeled as instantaneous events (i.e., CS onset), while the observational US/no US events were 1.5 seconds (i.e., from US onset to CS offset). There were only two types of events in the direct-expression stage, CS+ and CS-, modeled as 9 seconds (entire presentation of CS). Additionally, we included temporal modulation of the 1st order to capture effects of extinction in the direct-expression stage. In addition to the event regressors described above, we had six motion parameters (translation and rotation) as regressors of no interest. We added one delta regressor for each volume annotated by fMRIPrep as a motion outlier (up to 60 such volumes per subject in the observational learning stage, up to 38 in the direct-expression stage, median 8 and 2, respectively).

The US > no US was the primary contrast of interest in the observational learning stage and the CS+ > CS- in the direct-expression stage. We calculated the contrasts within-subject and used the parameter estimates in second-level analyses. We performed two types of second-level analyses in SPM: one-sample t-test designs for data pooled across groups (the ’friend’ and ’stranger’ groups) and two-sample t-tests for between-groups comparisons (the ’friend’ vs. the ’stranger’ group). Additionally, we used a temporal modulation regressor (CS+ × t) for both groups analyzed together and for between-group comparisons. We thresholded the second-level statistical maps using family-wise error (FWE) correction, with a p = 0.05 threshold. We used the voxel-level correction for the primary contrasts and cluster size correction with p = .001 cluster defining threshold for the results of temporal modulation and psychophysiological interactions.

#### 2.8.2. Region-of-interest - Definitions and Analysis

To perform the ROI analysis (parameter estimate extraction), the psychophysiological interaction analysis (time course extraction), and, in the case of the amygdala, the small volume correction (SVC) analysis, we defined anatomical regions of interest (ROI). We used the following brain structures as ROIs: the anterior insula (AI), anterior mid-cingulate cortex (aMCC), amygdala, right fusiform face area (rFFA), right posterior superior temporal sulcus (rpSTS), and right temporo-parietal junction (rTPJ). We chose regional definitions for these structures based on a meta-analytic connectivity mapping study focusing on social processes (Alcalá-López et al. 2018) and the Neurovault collection (https://identifiers.org/neurovault.collection:2462).

In the case of AI and amygdala, we combined anatomical masks from the left and right hemispheres and treated them as single ROIs. For the FFA, TPJ and pSTS, the regions considered functionally lateralized (Alcalá-López et al., 2018; Boccadoro et al., 2019; Sliwinska & Pitcher, 2018; Yovel et al., 2008), we used only the right-hemispheric masks. For the aMCC, which is a centrally-located structure, we used no hemispheric division. We used the same definitions across the ROI and PPI analyses. For small volume correction, which was applied only for the amygdala, we used an alternative bilateral definition based on the Harvard - Oxford atlas (Desikan et al., 2006) thresholded at 0.7. This definition was consistent with a previous study on observational fear conditioning (Lindström et al., 2018) and provided better structure coverage than the more restrictive meta-analytic definition. For the ROI analysis, the first-level parameter estimates were extracted and averaged within a given region for each participant using the spm_summarise function from SPM and compared using the BayesFactor package in R (Morey & Rouder, 2018).

#### 2.8.3. Psychophysiological Interactions

To perform the psychophysiological interaction analysis (PPI), we extracted the functional time series (1st eigenvariate) from the ROIs defined above and multiplied with the psychological variable (observation stage: US > no US; direct-expression stage: CS+ > CS-) using SPM VOI and PPI modules. We entered the PPI time series into a first-level model as the regressor of interest. We also included the functional time series and the psychological variable in the model as regressors and the motion parameters and motion outliers (the same as in the models used for activation analysis). Then we subjected parameter estimates obtained for the PPI regressor to second level analyses, using one-sample (both groups analyzed together) and two-sample (between-group) designs. We thresholded the statistical maps using cluster-based FWE correction, with cluster-defining threshold of p = 0.001 and corrected p-value of 0.05. Additionally, we applied a small volume correction to the amygdala.

#### 2.8.4. Statistical Analysis

To perform most statistical analyses, we used R with *afex*, *emmeans*, and *BayesFactor* libraries, with plots made using *the ggpubr* library. We calculated Bayesian ANOVA using JASP. We used repeated-measures ANOVA (with type III errors) and Bayesian repeated-measures ANOVA to analyze STAI-state scores (with the group as a between-subject factor and the questionnaire filling time, before and after the experiment, as a within-subject factor) and skin conductance response amplitudes (with the group and stimulus factors). We used the Wilcoxon-Mann-Whitney test to compare the discrete ratings describing the demonstrator’s expression (observational US) between the groups and a chi-squared test to compare the distribution of contingency-aware participants. We used t-test (with Welch’s correction for unequal variance) and Bayesian t-test (using default effect size priors, i.e., Cauchy scale 0.707) to compare the results of STAI-trait and subscales of SWE, and region-of-interest parameter estimates from fMRI.

For the classical tests, we applied a significance level of p = 0.05. For Bayesian t-tests, we used BF_10_ to denote evidence for the alternative hypothesis and BF_01_ for the null hypothesis. In Bayesian ANOVA, we reported the effects as the Bayes factor for the inclusion of a given effect (BF_incl_), calculated as a ratio between the models’ average likelihood, including the factor (given the data) vs. that of the models without the factor. When interpreting these results, we followed the convention suggested by Keysers et al. (2020) and considered ⅓ < BF < 3 as absence of evidence, 1/10 < BF < ⅓ or 3 < BF < 10 as moderate evidence and BF < 1/10 or BF > 10 as strong evidence.

## 3. Results

### 3.1. Behavioral Results

#### 3.1.1. Stimulus Contingency Awareness

After completing both stages of the experiment, the observers answered a set of questions about the relationship between the electrical stimulation and the visual cues in the observational learning stage (see section 2.4.4). 35 out of 48 participants in the ‘friend’ group and 35 out of 44 participants in the ‘stranger’ group correctly identified the CS+/US contingency. While the previous studies have reported rare cases of contingency-unaware participants (Golkar et al. 2015; Haaker et al. 2021), our ratio of the CS+/US contingency unaware subjects is higher. The high ratio is most likely a consequence of a real-time demonstrator-observer interaction, contributing to the less structured and thus more demanding learning context. A similar effect was observed in our previous study (Szczepanik, Kaźmierowska, et al., 2020). The proportion of contingency-aware and non-aware participants was not significantly different between groups (chi-squared = 0.554, df = 1, p = 0.46). We included only the contingency-aware participants in further analyses as we found that the learning efficacy depends on the contingency knowledge (Szczepanik, Kaźmierowska, et al., 2020).

#### 3.1.2. Evaluation of the Demonstrator’s Expression (the Observational US)

Next, the observers evaluated the demonstrator’s expression (the observational US) by answering a set of questions scored on 0 - 9 scales. The questions concerned the demonstrator’s reactions to electrical stimulation (how much discomfort they experienced, how expressive they were, and how natural their reactions were) and their attitude (how much empathy they felt for the demonstrator). Additionally, in the pairs of friends, the observers assessed how well they could identify with the demonstrator (this question was not asked in the ‘stranger’ group, see section 2.4.5.). Median ratings in both the ‘friend’ and ‘stranger’ groups were between 5 and 7 (see Figure 3). Additionally, we asked the observers to rate the degree of unpleasantness attributed to the demonstrators on a 1 - 5 scale (from very unpleasant to very pleasant). The median ratings were between 1 (very unpleasant) and 2 (rather unpleasant). The Wilcoxon-Mann-Whitney test showed no significant differences between the groups in any category. See Table S3 in the Appendix for detailed statistics.

**Figure 3:**
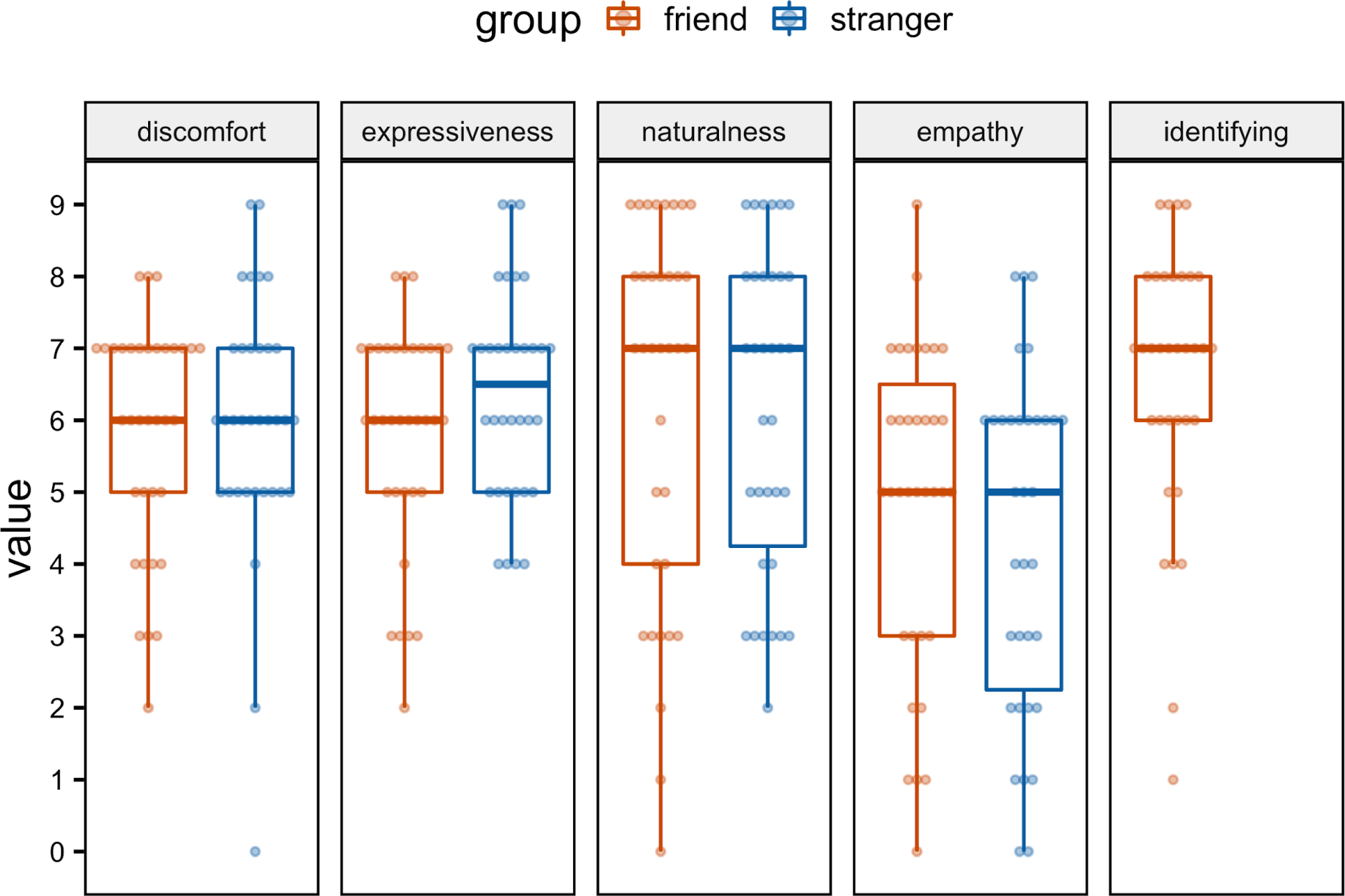
Evaluation of the Demonstrator’s Expression (the observational US) Note. Error bars extend to data points placed no further than 1.5*IQR (interquartile range) beyond the 1^st^ quartile and above the 3^rd^ quartile. The identification rating was done in the ‘friend’ group only.

#### 3.1.3. Empathy and Anxiety - Questionnaire Results

The state anxiety was tested twice, before and after the experiment. The results were analyzed for the contingency-aware observers, who were included into the fMRI analysis. Repeated-measures ANOVA and analogous Bayesian repeated measures ANOVA with the group (friend, stranger) as a between-subject factor and measurement (before, after) as a within-subject factor showed no significant differences: main effect of the group: F(1, 67) = 0.76, η_g_ ^2^ = .009, *p* = .39, BF = 0.35; main effect of the measurement: F(1,67) = 1.92, η_g_ ^2^ = .007, p = .17, BF_incl_ = 0.36; group × measurement interaction F(1,67) = 1.29, η_g_ ^2^ = .004, p = .259, BF_incl_ = 0.14.

The trait anxiety index, and three subscales of the empathic sensitiveness scale were treated independently, and compared between friend- and stranger-observers groups using two sample t-tests. Again, none of the comparisons was statistically significant and indicated either absence of evidence or moderate evidence against the difference: trait anxiety: t = 1.07, df = 63.5, p = 0.29, BF = 0.40; empathic concern: t = 0.20, df = 59.0, p = 0.85, BF = 0.25; personal distress t = 1.11, df = 64.1, p = 0.27, BF = 0.42; perspective taking: t = 0.12, df = 63.2, p = 0.91, BF = 0.25.

### 3.2. Skin Conductance Responses

Amplitudes of the observers’ skin conductance responses (SCR) were compared (separately for each stage of the experiment) using classical and Bayesian repeated measures ANOVA, with group (friend, stranger) as a between subjects factor. In the observational learning stage, ANOVA revealed a significant main effect of the stimulus (US / no US), F(1, 52) = 29.19, η_g_^2^ = .089, p < .001, BF_incl_ = 6976 where participants showed stronger reactions to the US compared to no US, t(52) = 5.40, p < .001, BF = 11341 (Figure 4 A). The main effect of the group, F(1,52) = 2.21, η_g_^2^ = .034, p = .14, BF_incl_ = 0.75, and the stimulus × group interaction, F(1,52) = 0.55, η_g_^2^ = .002, p = .46, BF_incl_ = 0.63, were not significant. Similarly, in the direct-expression stage, we found a significant effect of the stimulus (CS+/CS-), F(1, 52) = 5.73, η_g_^2^ = .018, p = .02, BF_incl_ = 1.79, with stronger reactions to CS+ than CS-, t(52) = 2.39, p = .02 , BF = 2.02 (Figure 4 B), while the main effect of the group, F(1, 52) = 1.89, η_g_ ^2^ = .029, p = .18, BF = 0.62, and the stimulus × group interaction, F(1,52) = 0.83, η_g_ ^2^ = .003, p = .37, BF = 0.464, were not significant.

**Figure 4:**
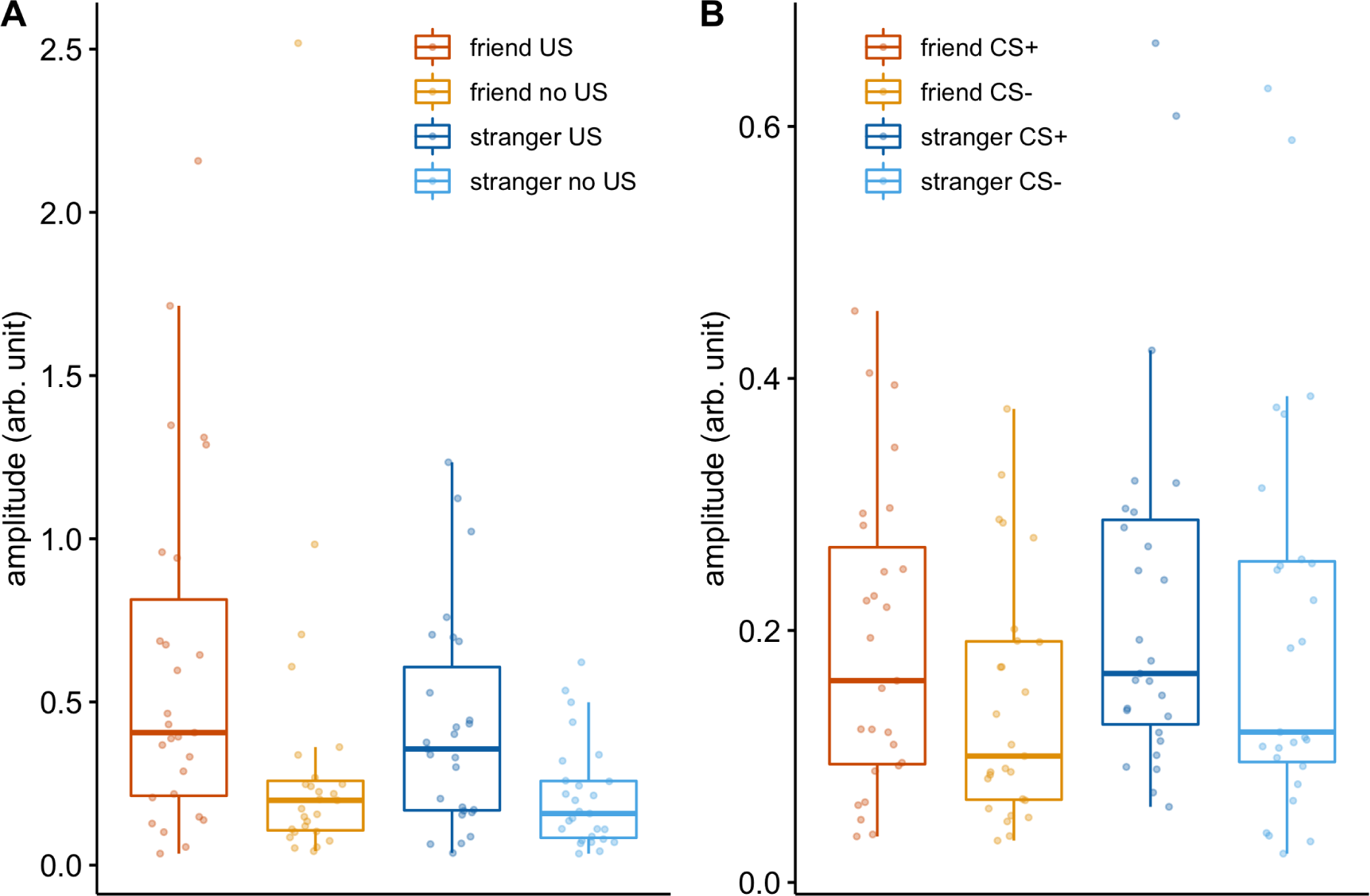
Skin Conductance Responses Note. (A) observational learning stage, (B) direct-expression stage. Error bars extend to data points placed no further than 1.5*IQR beyond the 1^st^ quartile and above the 3^rd^ quartile.

### 3.3. Imaging Results

#### 3.3.1. Brain Activation Analysis - Observational Learning Stage

In the observational learning stage, we analyzed the whole-brain activity of the observers witnessing the demonstrator’s reaction to the aversive stimulation. We compared it to the corresponding periods when the CS appeared without aversive stimulus (US > no US). To test the main effect of the task, we first evaluated the US > no US contrast for both groups. It revealed a robust and extensive activation (p < 0.05, FWE peak-level correction) of multiple brain regions, including the bilateral amygdala, bilateral anterior insula (AI), and anterior mid-cingulate cortex (aMCC), the bilateral posterior superior temporal sulcus (pSTS) and bilateral fusiform gyrus, including the fusiform face area (FFA).

Next, we compared the ‘friend’ and ‘stranger’ groups directly, i.e., with the (friend US > no US) vs. (stranger US > no US) using t contrasts (see Figure 5 A). The whole-brain analysis yielded no significant between-group differences (with either peak- or cluster-level FWE corrections). To further compare the groups, we performed region-of-interest analysis (averaging parameter estimates within a region) for the six pre-selected areas, defined independently of the functional data (AI, aMCC, amygdala, rFFA, rpSTS, rTPJ). We found no significant differences between the ‘friend’ and ‘stranger’ groups; Bayes Factor analysis indicated moderate evidence for the absence of effects (BF_01_ > 3) in all regions except the rFFA and rTPJ, where the evidence was inconclusive (BF_01_ = 2.5 and 2.07 respectively), see Table 1 and Figure 5 B.

**Figure 5:**
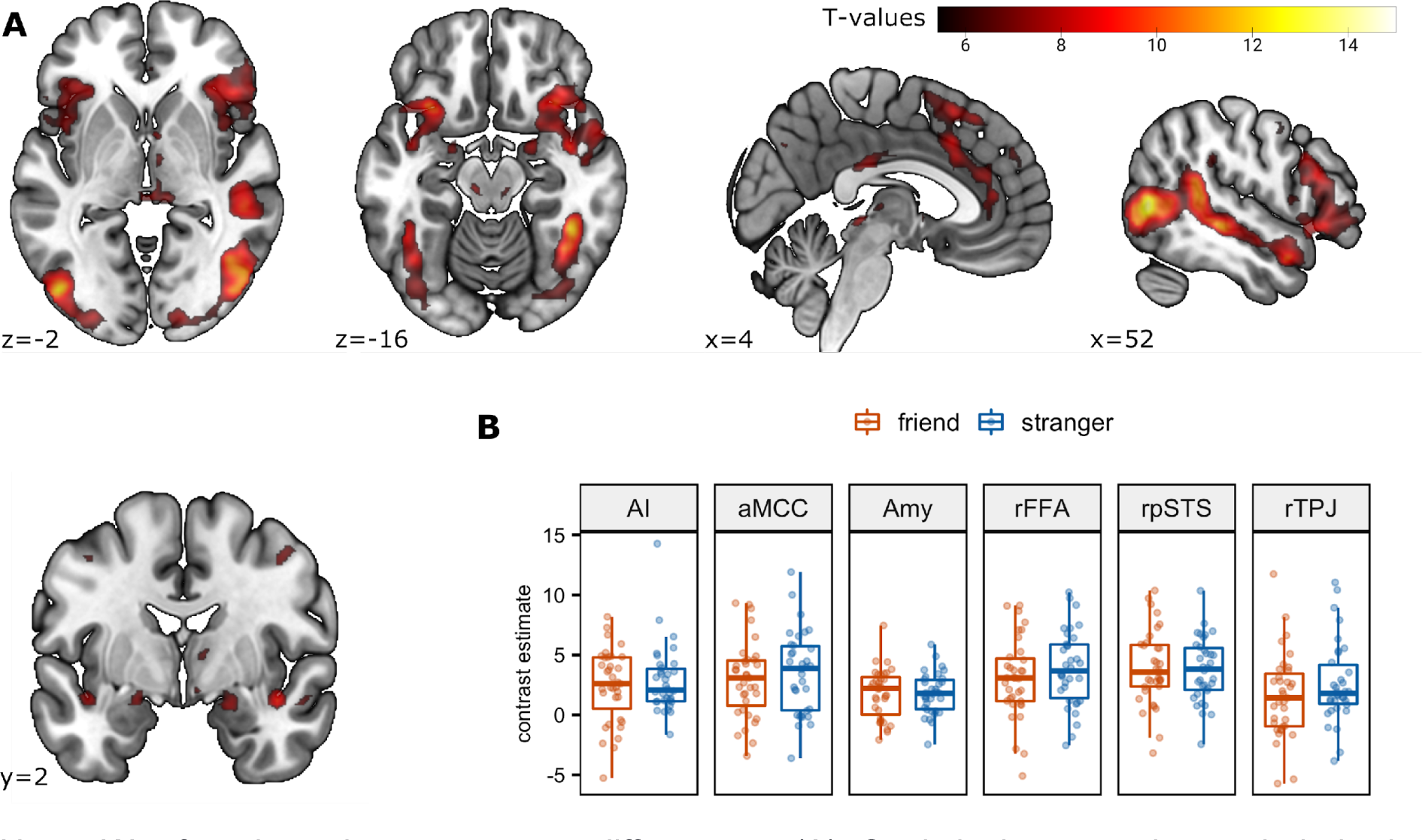
Activation in the Observational Learning Stage, US > no US Contrast Note. We found no between-group differences. (A) Statistical maps show whole-brain activations for both groups analyzed together, p < 0.05, FWE peak-level correction. (B) Region-of-interest statistics, showing group averages and 95% confidence intervals. Abbreviations: AI - anterior insula, aMCC - anterior mid-cingulate cortex, Amy - amygdala, rFFA - right fusiform face area, rpSTS - right posterior superior temporal sulcus, rTPJ - right temporo-parietal junction.

**Inline Supplementary Figure 5:**
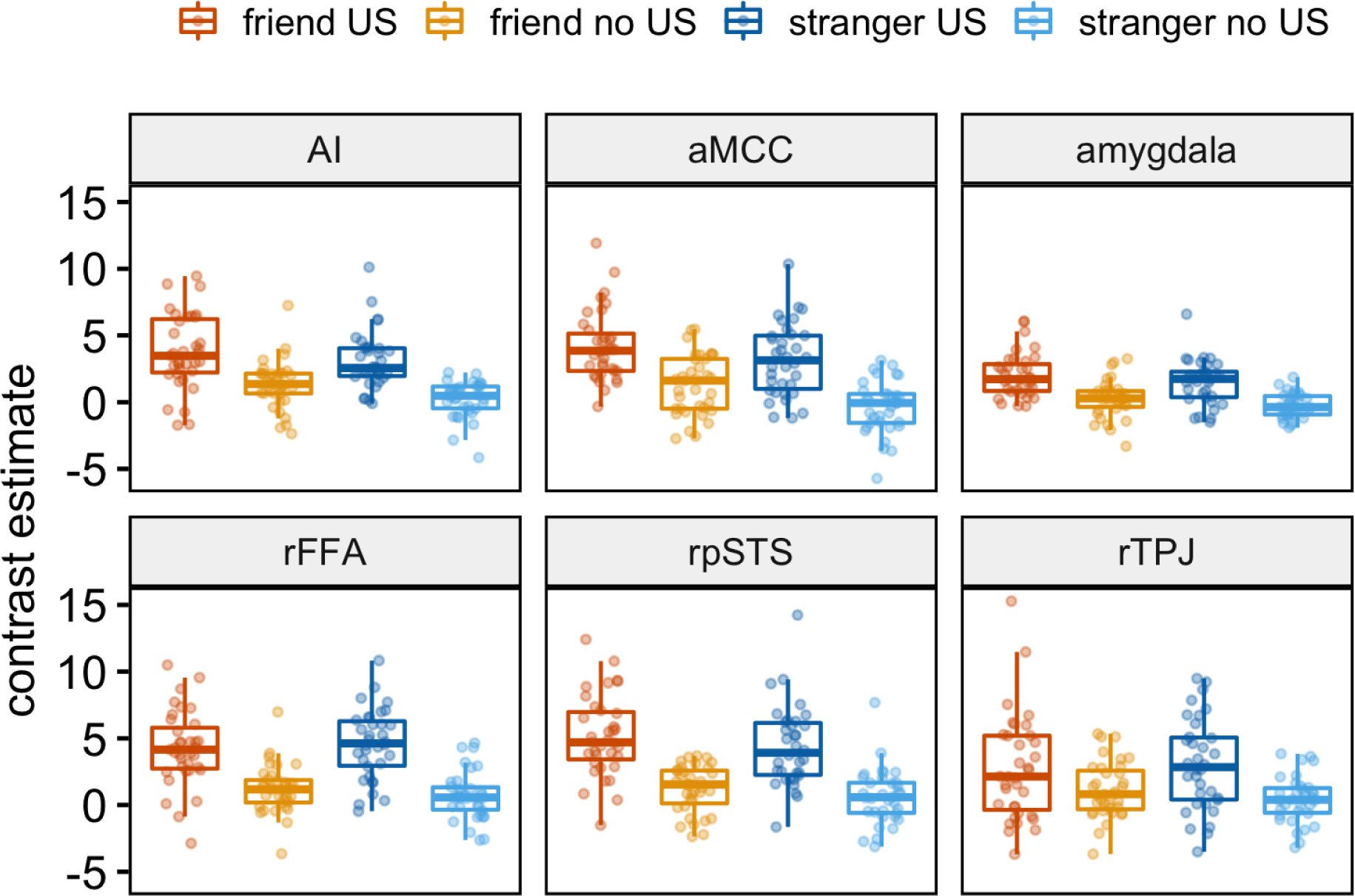
Region-of-interest Statistics for the Observational Learning Stage Note. Parameter estimates (group average and 95% confidence intervals, with subjects represented by dots) are shown for the US and no US separately. AI - anterior insula, aMCC - anterior mid-cingulate cortex, rFFA - right fusiform face area, rpSTS - right posterior superior temporal sulcus, rTPJ - right temporo-parietal junction.

**Table 1:** Region-of-interest Analysis of Between-group Differences

| ROI | Observational Learning Stage (US > noUS) |  |  | Observational Learning Stage (US) |  |  | Direct-expression Stage (CS+ > CS-) |  |  |
| --- | --- | --- | --- | --- | --- | --- | --- | --- | --- |
|  | t | p | BF<sub>01</sub> | t | p | BF<sub>01</sub> | t | p | BF<sub>01</sub> |
| AI | -0.64 | .52 | 3.39 | 1.06 | .29 | 2.50 | 0.03 | .97 | 4.04 |
| aMCC | -0.77 | .45 | 3.13 | 1.60 | .11 | 1.36 | -0.08 | .93 | 4.03 |
| Amy | 0.01 | .99 | 4.04 | 1.38 | .17 | 1.79 | 0.40 | .68 | 3.76 |
| rFFA | -1.06 | .29 | 2.50 | -0.49 | .62 | 3.65 | -1.08 | 0.29 | 2.47 |
| rpSTS | 0.24 | .81 | 3.94 | 1.08 | .29 | 2.47 | 1.14 | .26 | 2.32 |
| rTPJ | -1.25 | .21 | 2.07 | -0.40 | .69 | 3.78 | -0.13 | .90 | 4.01 |

*Note.* We performed two comparisons for the observational learning stage based on within-subject parameter estimates for either the (US > no US) or the US alone. We compared the (CS+ > CS-) contrast estimate between the groups for the direct-expression stage. The statistical results: positive t values represent friend > stranger; t-tests with Welch correction to the degrees of freedom; p values not corrected for the number of comparisons; Bayes factors reported in favor of the null hypothesis. AI - anterior insula, aMCC - anterior mid-cingulate cortex, Amy - amygdala, rFFA - right fusiform face area, rpSTS - right posterior superior temporal sulcus, rTPJ - right temporo-parietal junction.

An additional between-group comparison, based on the observational US reactions, i.e., friend US > stranger US t contrast, also did not yield significant differences on a whole-brain level. Region-of-interest analysis for this contrast (using the same set of regions as above) confirmed the lack of significant between-group differences. At the same time, Bayes factors indicated evidence that was either inconclusive or moderately favored the null hypothesis (all BF_01_ between 1.36 and 3.65), see Table 1 and Inline Supplementary Figure 5.

#### 3.3.2. Brain Activation Analysis - Direct-expression Stage

To measure observational fear learning, i.e., the association formed between the CS and the observational US, in the direct-expression stage, we analyzed the brain activity of the observers in response to the conditioned stimuli (CS+ > CS-). To evaluate the main effect of the task, the CS+ > CS- contrast was first analyzed for both groups together and subsequently compared between the groups.

Analyzing both groups together, we observed significant activations in the bilateral AI and the aMCC (Figure 6). Next, we performed a direct between-group comparison, i.e., friend (CS+ > CS-) vs. stranger (CS+ > CS-), which yielded no significant differences on the whole-brain level. We performed a region-of-interest analysis with the same set of regions as in the observational learning stage to further compare the groups. We found no significant differences, and Bayes Factor analysis indicated moderate evidence for the absence of effects (BF_01_ > 3) in all regions except the rFFA and rpSTS where the evidence was inconclusive (BF_01_ = 2.47 and 2.32 respectively); see Table 1 and Figure 8.

**Figure 6:**
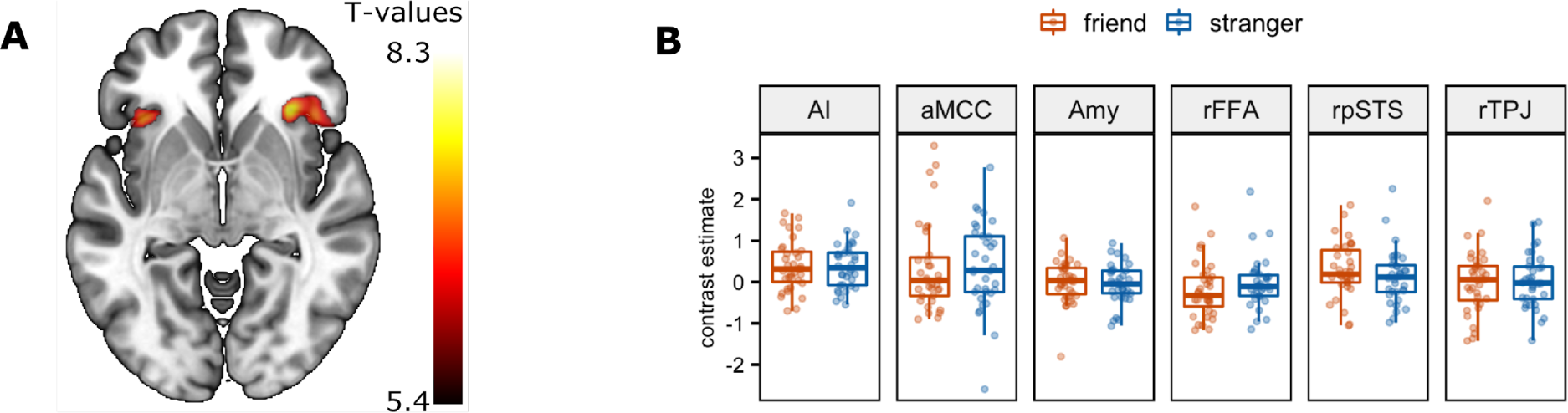
Activation in the Direct-expression Stage, CS+ > CS- Contrast Note. (A) Statistical map showing whole-brain activations for the ‘friend’ and ‘stranger’ groups analyzed together, FWE corrected, p < 0.05. (B) Region-of-interest statistics, showing group averages and 95% confidence intervals. Abbreviations: AI - anterior insula, aMCC - anterior mid-cingulate cortex, Amy - amygdala, rFFA - right fusiform face area, rpSTS - right posterior superior temporal sulcus, rTPJ - right temporo-parietal junction.

To capture the potential effects of extinction during the direct-expression stage, we included 1st order temporal modulation in the statistical model. Analyzing both groups together, we observed a significant decrease in the brain responses to CS+ across several regions, most notably in the aMCC. To test whether the groups differed concerning the temporal dynamics of brain responses, we evaluated the (friend CS+ × t) vs. (stranger CS+ x t) contrast. We observed the differences in some brain regions, including the left inferior frontal gyrus, left middle temporal gyrus, right lingual gyrus, and left putamen. Strangers showed a stronger linear decrease in activity than friends (see Figure 7 and Table S6).

**Figure 7:**
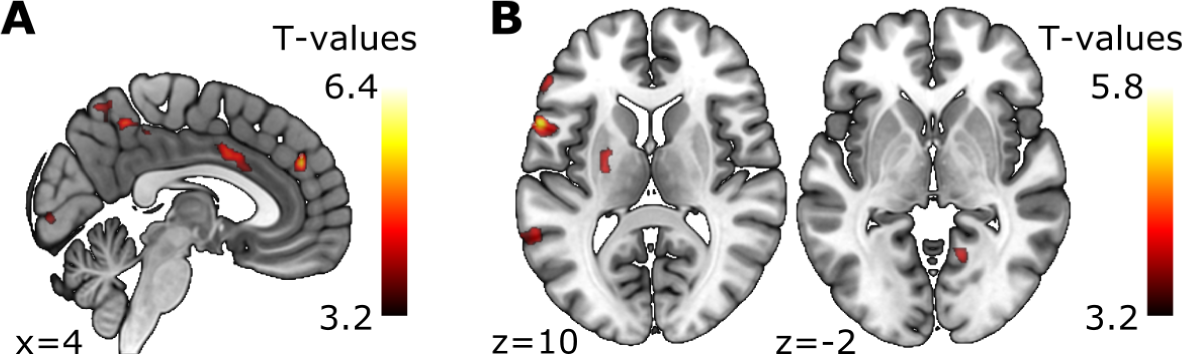
Temporal Modulation of the CS+ Response in the Direct-expression Stage Note. (A) Statistical map showing a significant effect for the ‘friend’ and ‘stranger’ groups analyzed together. (B) Statistical map showing the (stranger CS+ × time decrease) > (friend CS+ × time decrease) contrast.

Importantly, however, we found no differences in the structures activated in the observational and direct-expression stages.

#### 3.3.3. Psychophysiological Interactions

We performed a psychophysiological interaction (PPI) analysis for several selected ROIs to investigate the coupling between the activated structures. First, we analyzed the observational learning stage, focusing on the US > no US contrast. As the seed structures, we used the AI, rpSTS, aMCC, amygdala, rFFA, and rTPJ. Analyzing both groups together, we observed increased coupling of the AI with several regions, including the rpSTS (Figure 8 A), and stronger coupling of the rpSTS with the AI, right fusiform gyrus (Figure 8 B), and amygdala (Figure 8 C). However, there were no significant differences between the groups. Similarly, we did not find the group differences using the aMCC, amygdala, rFFA, and rTPJ as the seed structures. Further, we applied the PPI analysis to the activations observed during the direct-expression stage, using the AI, aMCC, and amygdala as the seeds. We did not find significant differences for the groups analyzed together or between the friend and stranger groups.

**Figure 8:**
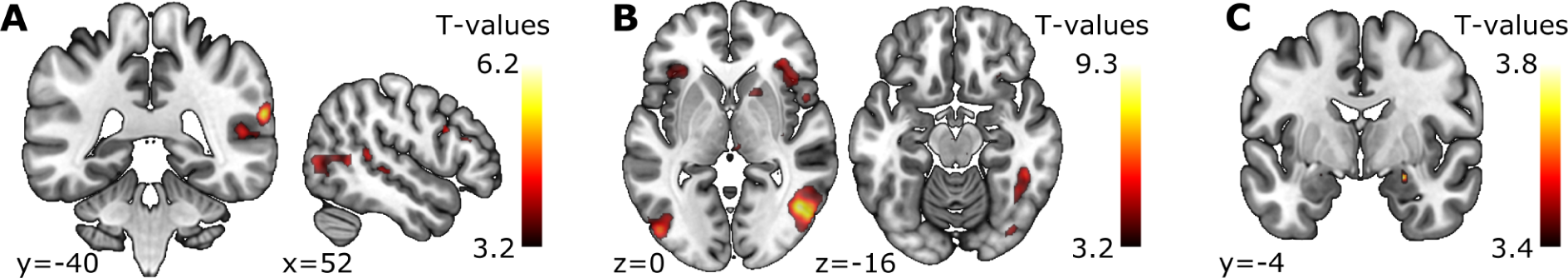
Psychophysiological Interaction Maps, US > no US Contrast Note. AI and pSTS seeds were analyzed for both groups together during the observational learning stage. (A) the AI exhibits interaction with several regions, including the rpSTS (FWEc), (B) the rpSTS exhibits interaction with several areas, including the right fusiform gyrus and bilateral AI (FWEc), (C) psychophysiological interaction of the rpSTS with the amygdala (SVC).

#### 3.3.4. Multivariate pattern analysis

To explore if possible group differences could be encoded in the patterns of activations rather than in the activation strength in specific regions, we have tried a multivariate approach. We used classification (decoding), one of the most popular multivariate pattern analysis (MVPA) methods (Haynes 2015; Valente et al. 2021), and employed The Decoding Toolbox (Hebart et al. 2014). Due to the limitations of our experimental design (which was not planned for the MVPA analysis), we treat this analysis as exploratory and report it in the Appendix (see section 1). We found no statistically significant group differences in the classifier’s accuracy, neither in the observational fear learning nor in the direct-expression stage, which is consistent with the results of other analyses described in the manuscript.

## 4. Discussion

Here, we investigated the relevance of the demonstrator’s familiarity in observational fear learning using physiological and neural measures. We focused on the US > no US contrast in the observational learning and the CS+ > CS- contrast in the direct-expression stage. We found no significant differences between participants observing friends (‘friend’ group) and subjects watching strangers (‘stranger’ group), neither in fear contagion nor in observational fear learning. Our study is the first to directly examine the difference between neural correlates of observational fear learning from friends and strangers. Using the ecologically valid paradigm that has been developed and tailored for this study, we replicated the previously reported brain activations underlying fear conditioning in humans. However, brain activity did not differ between participants observing friends and strangers. This clarifies the role of familiarity in fear contagion in humans and is in line with the previous rodent reports (Hernandez-Lallement et al., 2020). Overall, our results show that humans learn fear equally from friends and strangers, which might have been evolutionarily beneficial.

### 4.1. Main Physiological and fMRI Effects - Validation of the Protocol

We improved the original protocol of Haaker, Golkar et al. (2017) to study a real-time demonstrator-observer interaction. To validate the modified protocol, we firstly examined the data regardless of the level of familiarity. We analyzed both physiological and neural correlates of observational fear learning. The majority of observers correctly identified the CS+/US contingency and the contingency-aware to non-aware participants ratio, around three-quarters, was similar across groups. As our previous study showed that the psychophysiological effects of observational fear learning were present only in the contingency-aware participants (Szczepanik, Kaźmierowska, et al., 2020), we included only those in the analyses.

We observed enhanced observers’ skin conductance response (SCR) to the shock-associated stimuli during both observational learning and direct-expression stages, which is in line with earlier reports (Sevenster et al., 2014; Szczepanik, Kaźmierowska, et al., 2020). Neuroimaging analysis of data collected during the observational fear learning stage (US > no US contrast) revealed brain activity in multiple regions, including those crucial for fear-related processing (amygdala, AI, aMCC), which is consistent with previous studies (Lindström et al., 2018). Additionally, we observed activation of brain areas involved in dynamic social perception (fusiform gyrus, pSTS, and supplementary motor area), which may indicate processing of visual body cues (Allison et al., 2000; Yang et al., 2015). PPI analysis revealed coupling of the AI with the rpSTS and of the rpSTS with the AI, right fusiform gyrus, and amygdala, suggesting that these brain regions are jointly involved in the social transfer of fear. Brain activity during the direct-expression stage (CS+ > CS- contrast) involved the bilateral insular cortex and aMCC, which is in line with previous findings (Lindström et al., 2018; Olsson et al., 2007). We did not observe significant amygdala activations. In the direct fear conditioning, the amygdala is considered a region where the CS-US association is stored (Olsson & Phelps, 2007; Phelps & LeDoux, 2005). However, none of the previous studies has provided consistent evidence for the amygdala involvement during the direct-expression stage. Additionally, a meta-analysis on the neural signatures of the human fear conditioning has shown that the amygdala is not reliably activated when fMRI fear conditioning tasks are employed (Fullana et al., 2016). Taken together, using the ecologically valid paradigm, we replicated most of the previously reported brain activations underlying observational fear conditioning in humans.

### 4.2. Familiarity Effects

We employed the experimental design in which familiarity with the demonstrator was the only between-group difference. We also made sure that the observers learned about the threat through social means only - they did not directly experience any electrical stimulation at any point of the experiment. The questionnaire results indicated that the observers from both groups did not differ in their general level of anxiety and empathy-related traits. Also, their initial state of anxiety (measured at the beginning of the experiment) was similar. Notably, the expressions of the demonstrators (the observational US) were rated as natural and did not differ between the groups. This lack of differences indicates that the experimental conditions were similar across groups, which is essential considering the naturalistic paradigm employed and potential confounds related to individual differences in the demonstrators’ fear expression. We found no significant differences in the brain activations of observers learning about the threat from their friends or strangers. The only previous report directly comparing the neural correlates of threat-to-self, threat-to-friend, and threat-to-stranger experience (Beckes et al., 2013) used a within-subject design. The within-subject protocol makes it difficult to disentangle the impact of direct vs. social sources of information. Our results show that familiarity between the participants does not modulate social fear learning when a demonstrator is a sole source of information.

In contrast to our results, the social learning of pain activated the pain network components (ACC and AI) depending on whether the participants imagined a loved one or a stranger (Cheng et al., 2010). This result seems to be at odds with our findings. However, the fear and pain paradigms differ. Although both paradigms use aversive stimulation (often involving electric shocks), the former emphasizes that it should be uncomfortable but not painful. In our study, we adjusted the electric shock’s intensity by asking every demonstrator to choose the level of electric stimulation and instructing him to react ‘in a natural, yet noticeable manner’. Based on the observers’ post-experimental ratings (on average describing the degree of unpleasantness attributed to the demonstrators as ‘rather unpleasant’), we may suppose they did not interpret the demonstrators’ condition in terms of pain.

Animal studies provide somewhat disparate results on the role of familiarity in threat contagion (Gonzalez-Liencres et al., 2014; Jeon et al., 2010; Knapska et al., 2010; Sanders et al., 2013). The studies carried out in a well-known environment, in which the presence of a partner attenuates threat response (the effect known as social buffering (Kiyokawa et al., 2014) more consistently report familiarity effect. Thus, familiarity may play a role in social attenuation rather than the social enhancement of fear. What is more, a recent meta-analysis of rodent studies indicated no familiarity effect on emotional contagion of threat (Hernandez-Lallement et al., 2020). The lack of familiarity effect in threat contagion may stem from the fact that social learning about danger, especially in a novel environment, is equally effective and might have been evolutionary beneficial regardless of the demonstrator’s level of familiarity.

Emotional contagion is considered a primary form of empathy, which results from the basic mechanism of the Perception-Action Model (De Waal, 2012; de Waal & Preston, 2017). The PAM involves automatic matching between the target’s and the observer’s neural responses and constitutes a foundation for more complex phenomena such as sympathetic concern and perspective-taking. The model predicts that emotional contagion is more robust among individuals in close social relationships who share past experiences as their perception-action coupling is stronger (Preston & de Waal, 2002). However, our data show that fear contagion does not depend on the level of familiarity, which may stem from its crucial informative role. Preston and de Waal’s model also postulates that emotional contagion is stronger when empathy is high. The reported level of experienced empathy toward the demonstrator did not differ between the groups, which could explain the lack of between-group differences in observational fear learning. Although surprising, a similar (and not exceptionally high) level of experienced empathy in both groups suggests the priority of threat processing over empathy involvement in our experimental situation. In the face of threat, observing the emotional responses of others can facilitate life-saving responses (Olsson et al., 2020). It is noteworthy that the actors serving as demonstrators in previous observational fear learning studies were all unfamiliar to the observers and still remained an effective source of learning (Golkar et al., 2015; Golkar & Olsson, 2017). Thus, perceiving the emotions of others not only provides information about their source, i.e., the emotional state of the ‘demonstrator’ but also about the environment. Friends and strangers are equally good sources of information about threats.

### 4.3. Limitations

Since the participants came together to the laboratory in the ‘friend’ group, the situation was more natural and socially engaging than in the ‘stranger’ group, where the observers had no interaction with the demonstrators before the experiment. However, if this was an essential factor, we could expect stronger activations of the social brain network in the ‘friend’ group, which was not the case. Moreover, we did not directly measure observers’ empathy toward the demonstrators. However, we assessed observers’ general level of empathy, situational empathy experienced during the observation and degree of unpleasantness attributed to the demonstrator and did not find differences between the groups. Finally, we studied only males; thus, it is unclear whether the results extrapolate to the female population. Considering the sex differences in emotional processing and empathy (Proverbio, 2021) and a pioneering character of the study, limiting the probe to one sex only enabled us to watch relatively strong and statistically powered effects. However, this issue certainly requires future investigations.

### 4.4. Conclusions

We describe here the neural correlates of observational fear learning in humans using a naturalistic approach. Using this paradigm, we replicated most of the previously reported brain activation underlying fear conditioning in humans. Importantly, our findings constitute the first neuroimaging evidence for the lack of relationship between familiarity and fear contagion and observational fear learning in humans. We argue that fear contagion is an automatic, unconscious process independent of the level of the demonstrator’s familiarity. We claim that fear is learned equally from friends and strangers in humans, which is evolutionarily beneficial. These results resonate with ongoing debates on the questions such as the components of empathy, the boundaries between unconscious threat processing and conscious fear experience, and the role of the amygdala in the fear conditioning process. A question of the demonstrator’s characteristics modulating the observational fear learning process remains open, creating a space for further investigations.

## Data Availability

Relevant data are stored in an OSF repository and are available at https://osf.io/g3wkq/. Unthresholded statistical maps from the reported comparisons are also available at Neurovault, https://neurovault.org/collections/RSLLSFTQ/. Code replicating analyses reported here is available at https://github.com/nencki-lobi/emocon-mri.

## Acknowledgements

We thank Jan Haaker for his valuable advice regarding the protocol’s modifications. We also acknowledge Urszula Baranowska and Małgorzata Dąbkowska for their support in collecting data, as well as Michał Kaźmierowski, Monika Riegel and Małgorzata Wierzba for their helpful insights regarding data analysis. Data collection and analysis were sponsored by National Science Centre grant 2015/19/B/HS6/02209. Ewelina Knapska was supported by European Research Council Starting Grant (H 415148).

## Author Contributions

Anna M. Kaźmierowska - Conceptualization, Methodology, Data curation, Formal Analysis, Investigation, Software, Visualization, Writing - original draft, Writing - review & editing

Michał Szczepanik - Conceptualization, Methodology, Data curation, Formal Analysis, Investigation, Software, Visualization, Writing - original draft, Writing - review & editing

Marek Wypych - Conceptualization, Methodology, Supervision, Writing - review & editing Dawid Droździel - Investigation, Resources

Artur Marchewka - Conceptualization, Methodology, Resources, Supervision, Writing - review & editing

Jarosław M. Michałowski - Conceptualization, Methodology, Supervision, Writing - review & editing

Andreas Olsson - Conceptualization, Supervision, Writing - review & editing

Ewelina Knapska - Conceptualization, Methodology, Funding acquisition, Project administration, Supervision, Writing - review & editing

