## Appendix for "Learning about threat from friends and strangers is equally effective: an fMRI study on observational fear conditioning"

### 1. MVPA analysis

To explore if possible group differences could be encoded in the patterns of activations rather than in the strength of activation in specific regions, we have tried a multivariate approach. We used classification (decoding), one of the most popular multivariate pattern analysis (MVPA) methods (Haynes, 2015; Valente et al., 2021) and employed The Decoding Toolbox (Hebart et al., 2014).

Because of the limitations of our experimental design (which was not planned for the MVPA analysis; i.e., had only one run per task, containing 12 trials of each type), we carried out an inter-subject pattern analysis (Wang et al., 2020), in which training and testing was done across subjects, rather than across runs or trials within each participant. We used the leave-one-subject-out cross-validation scheme, and analyzed friend and stranger groups separately.

We performed the classification within six regions of interest (ROI). We used the same anatomical masks as in the univariate ROI analyses: the bilateral amygdala (AMY), bilateral anterior insula (AI), anterior mid-cingulate cortex (aMCC), right fusiform face area (rFFA), right posterior superior temporal sulcus (rpSTS) and right temporo-parietal junction (rTPJ). The classification accuracy was calculated for each ROI separately.

We used beta files from first-level models as the input for the classification for each subject. The single-subject models used the images preprocessed by fMRIPrep (as described in the main text), but more subtle smoothing was applied for MVPA (2 mm FWHM 3D Gaussian kernel). While unsmoothed images are often used in multivariate analyses, a small amount of smoothing contributes to noise reduction, which improves decoding accuracy (Wang et al., 2020). The conditions of interest were the US and no US in the observational learning stage and CS+ and CS- in the direct-expression stage.

The output measures for each group were the ‘accuracy minus chance’ values (one per group). The accuracy was calculated as the mean percentage of correct classifications (of both the US and no US in the learning stage or CS+ and CS- in the direct-expression stage) averaged across n cross-validation folds. We subtracted the chance level (50%) from the accuracy values and then computed the difference between the groups.

We further tested the obtained between-group differences (one per each ROI and each task) against the distribution of 1000 analogous chance differences. To do this we randomly divided subjects into two groups (equal in size to original friends and strangers groups) and applied the procedure described above to compute the between-group classification difference. This was repeated 1000 times, and resulted in a 'null' distribution of 1000 differences between the randomly divided groups. The actual group differences presented against the null distributions are presented in Figures S1 & S2. The p-values were calculated as a percentage of the differences whose absolute values exceeded the absolute value of the actual group difference. The between-group differences observed at both stages of the experiment and their corresponding p-values are shown in Table S1.

The described analysis did not reveal statistically significant group differences in the classifier's pattern recognition accuracy. However, a trend-level difference was found in the amygdala in the direct-expression stage, suggesting a higher classification accuracy of CS+ and CS- in the stranger- compared to friend-observers. This difference was driven by very poor (significantly below the mean for randomly divided groups) classification accuracy in the friends group and relatively high classification accuracy in the strangers group (Figure S3).

Due to the suboptimal design of our study, we treat the described analysis as exploratory and are cautious in formulating interpretations. Our attempt has been the first of this kind in observational fear learning, and it highlights the possible advantages of employing MVPA methods in further studies on this topic.

We run the classification analysis using The Decoding Toolbox (Hebart et al., 2014) with (default) SVM classifier. The decoding part was calculated based on a built-in template script. The code for calculating the decoding accuracy between the permutated subsets of data can be found at
https://github.com/nencki-lobi/emocon-mri.

**Figure S1**

*The US / no US Classification Results in the Observational Learning Stage*

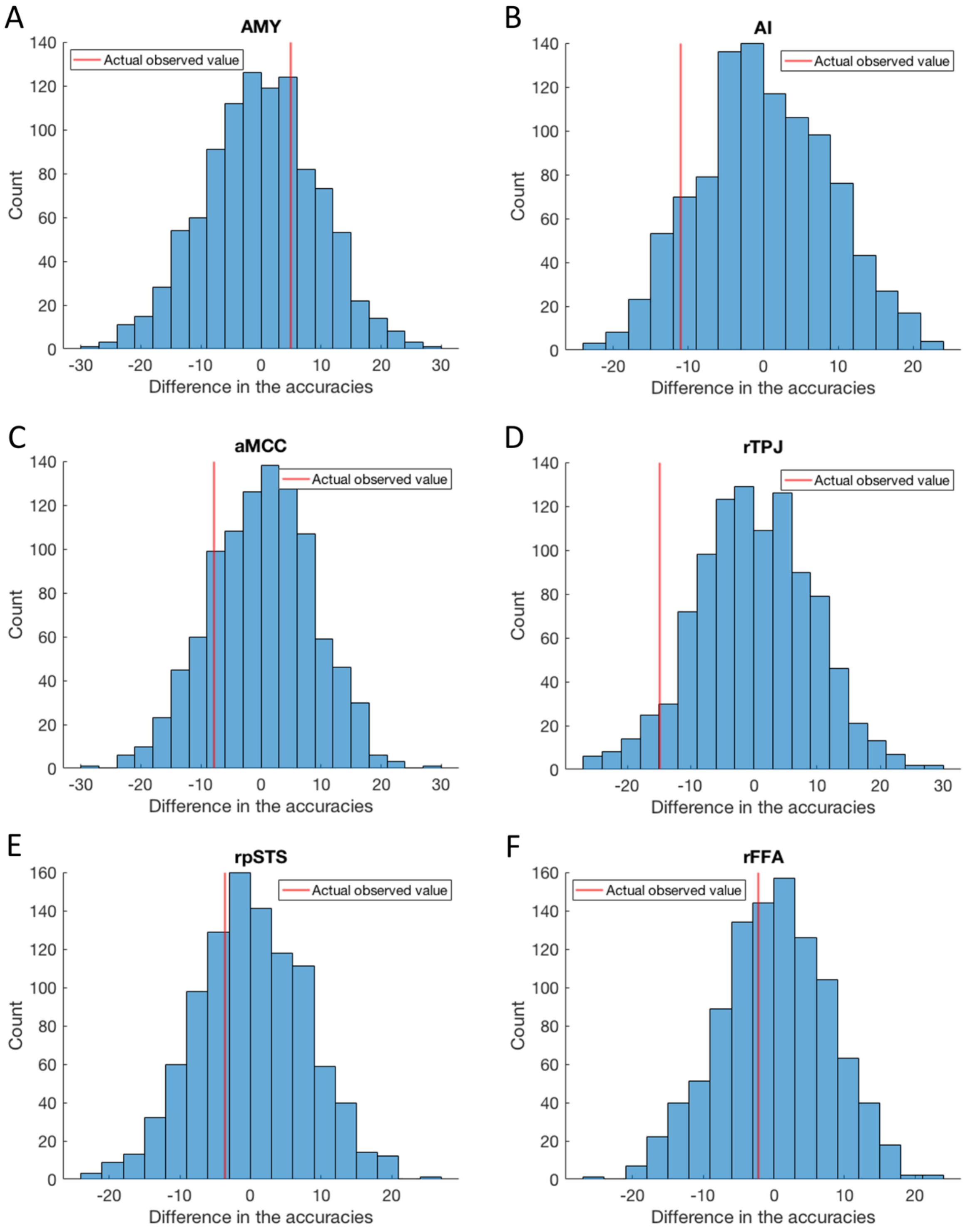

*Note.* The histograms show distributions of 1000 differences in accuracy between the randomly divided groups. The red line indicates a difference between the accuracies in the actual friends and strangers groups. (A) bilateral amygdala, (B) bilateral anterior insula, (C) anterior mid-cingulate cortex, (D) right temporo-parietal junction, (E) right posterior superior temporal sulcus, (F) right fusiform face area.

**Figure S2**

*The CS+ / CS- classification results in the Direct-expression Stage*

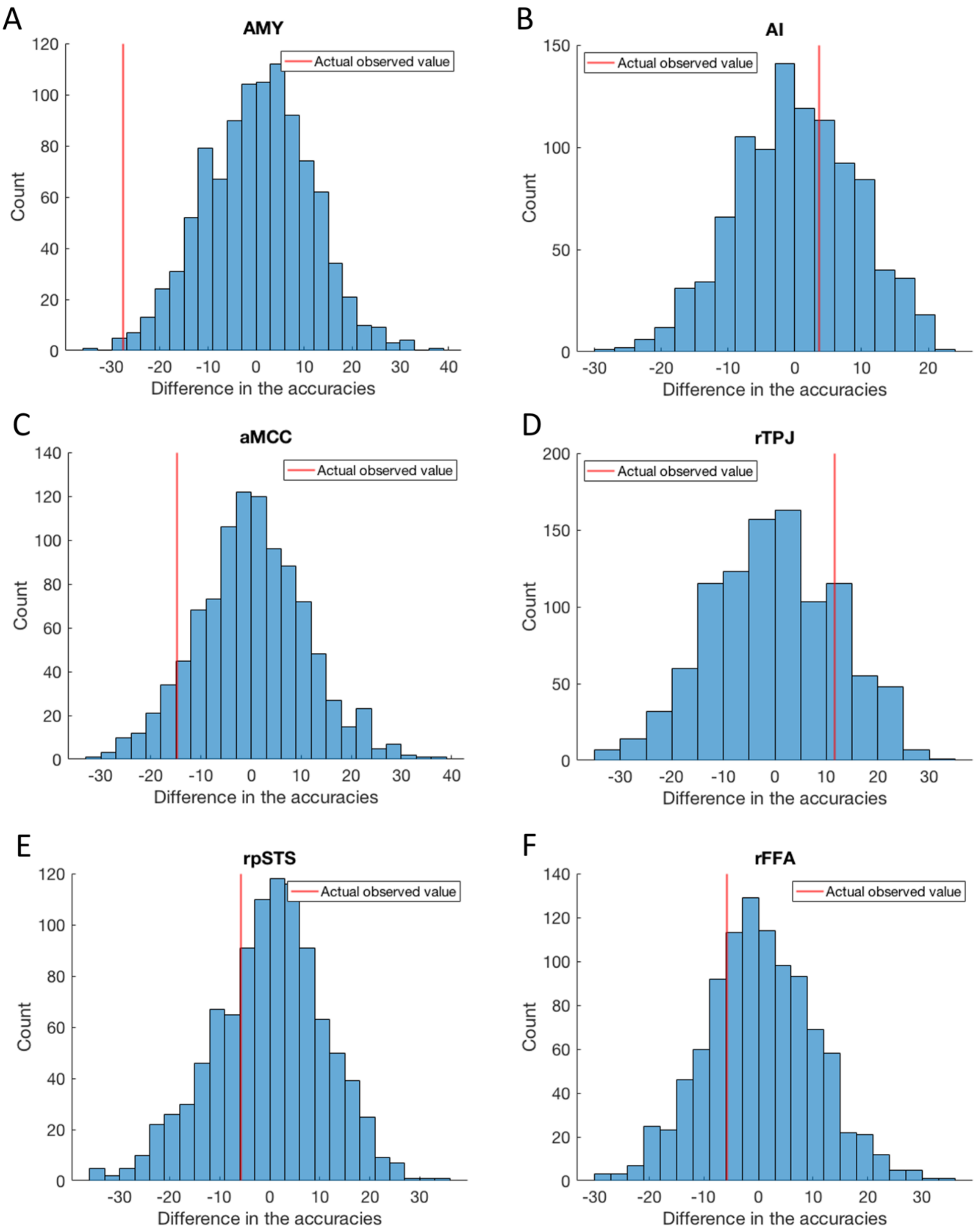


*Note.* The histograms show distributions of 1000 differences in accuracy between the randomly divided groups. The red line indicates a difference between the accuracies in the actual friends and strangers groups. (A) bilateral amygdala, (B) bilateral anterior insula, (C)

anterior mid-cingulate cortex, (D) right temporo-parietal junction, (E) right posterior superior temporal sulcus, (F) right fusiform face area.

**Table S1**

*The 'Accuracy Minus Chance' Values for Both Groups in Each ROI*

| ROI | Observational Learning Stage | | | | Direct-expression Stage | | | |
|---|---|---|---|---|---|---|---|---|
| | F | S | diff F-S | p | F | S | diff F-S | p |
| Amy | 30.00 | 25.00 | 5.00 | 0.60 | -5.71 | 22.06 | -27.77 | 0.07† |
| AI | 24.29 | 35.29 | -11.01 | 0.19 | 25.71 | 22.06 | 3.66 | 0.67 |
| aMCC | 14.29 | 22.06 | -7.77 | 0.38 | 1.43 | 16.18 | -14.75 | 0.17 |
| rTPJ | 7.14 | 22.06 | -14.92 | 0.11 | 7.14 | -4.41 | 11.55 | 0.37 |
| rpSTS | 22.86 | 26.47 | -3.61 | 0.66 | -5.71 | 0.00 | -5.71 | 0.63 |
| rFFA | 24.29 | 26.47 | -2.18 | 0.78 | -4.29 | 1.47 | -5.76 | 0.60 |

*Note.* The US and no US patterns were analyzed in the observational learning stage, while in the direct-expression stage, the CS+ and CS- patterns. A difference of the 'accuracy minus chance' values between the friends and strangers group was calculated for each ROI (column diff F-S). The result was tested against 1000 differences between randomly divided groups. The p-values of these tests (two-tailed) are presented in column p.

F - friends, S - strangers, Amy - the bilateral amygdala, AI - bilateral anterior insula, aMCC - anterior mid-cingulate cortex, rTPJ - right temporo-parietal junction, rpSTS - right posterior superior temporal sulcus, rFFA - right fusiform face area.

†Bonferroni-Holm correction for six comparisons applied.

**Figure S3**

*The CS+ / CS- Classification Results in the Direct-expression Stage in the Amygdala*

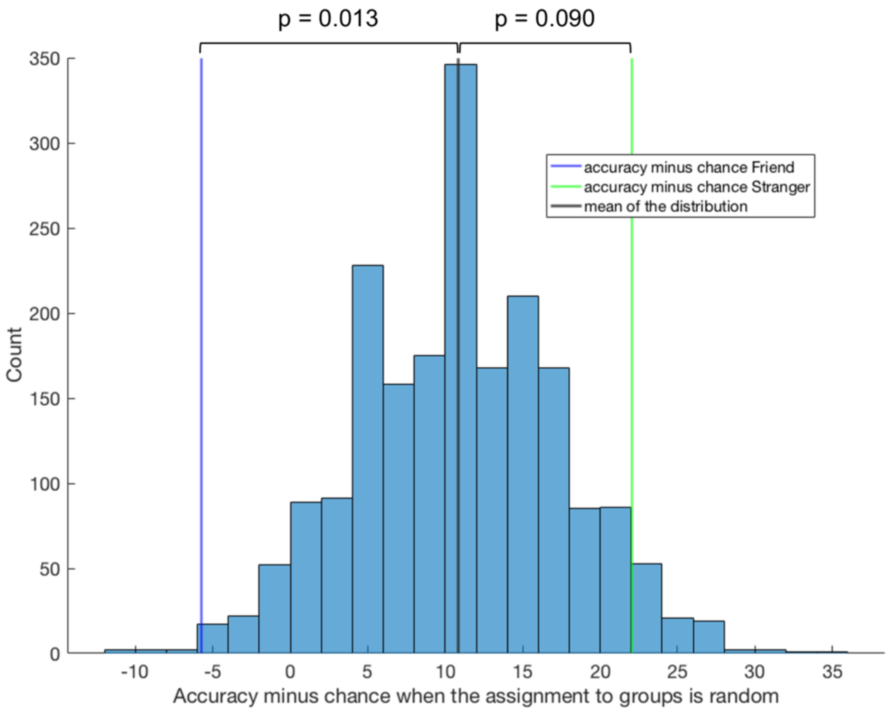


*Note.* The histogram shows 2000 'accuracy minus chance' values derived from two groups of 1000 random group divisions. The blue line marks the 'accuracy minus chance' value derived from the true friends group and the green line marks the 'accuracy minus chance' value derived from the true strangers group. The black line indicates the mean of the random distribution. p values (two-tailed) for two comparisons are provided.

### 2. Questionnaire results

**Table S2**

*Descriptive Statistics for Age, Demonstrator-observer Friendship Length and the McGill Friendship Questionnaire (MFQ) Result for the Subgroups of Participants*

| Variable | Subgroup | Mean (SD) | Median (IQR) | Range |
|---|---|---|---|---|
| Age | OF | 22.5 (2.77) | 22 (4.5) | 18 - 29 |
| | OS | 23.3 (2.95) | 22.5 (4) | 18 - 30 |
| | D | 22.1 (2.53) | 22 (4) | 18 - 28 |
| Friendship Length | OF | 7.7 (4.22) | 6 (7) | 3 - 19 |
| MFQ Result | OF | 52 (7.7) | 55 (9.5) | 33 - 60 |
| | DF | 49.8 (9.21) | 52.5 (15) | 30 - 60 |

*Note.* Data regarding friendship length and the MFQ result were collected in the ‘friend’ group only. OF: observers from the ‘friend’ group who learned the contingency (n = 35); OS: observers from the ‘stranger’ group who learned the contingency (n = 34); D: demonstrators for whom either a friend or a stranger observer learned the contingency (n = 44); DF: demonstrators for whom a friend observer learned the contingency (n = 34); SD: standard deviation; IQR: interquartile range.

**Table S3**

*Evaluation of the Demonstrator’s Expression (the Observational US)*

| | Discomfort of the Demonstrator | | Expressiveness of the Demonstrator | | Naturalness of the Demonstrator | | Empathy Toward the Demonstrator | | Identifying with the Demonstrator | | Unpleasantness Attributed to the Demonstrator | |
|---|---|---|---|---|---|---|---|---|---|---|---|---|
| Group | F | S | F | S | F | S | F | S | F | S | F | S |
| Median (IQR) | 6 (2.0) | 6 (2.0) | 6 (2.0) | 6.5 (2.0) | 7 (4.0) | 7 (3.75) | 5 (3.5) | 5 (3.75) | 7 (2.0) | n/a * | 1 (1.0) | 2 (1.0) |
| Range | 2 - 8 | 0 - 9 | 2 - 8 | 4 - 9 | 0 - 9 | 2 - 9 | 0 - 9 | 0 - 8 | 1 - 9 | n/a | 1 - 4 | 1 - 5 |
| Stat. Test | W = 581.5, p = 0.87 | | W = 501, p = 0.25 | | W = 630.5, p = 0.67 | | W = 671.5, p = 0.36 | | n/a | | W = 586, p = 0.91 | |

*Note.* To evaluate the demonstrator's expression (the observational US), we asked the observers to rate (on a 0 - 9 scale) the demonstrator's reactions. The questions concerned the demonstrator's responses to electrical stimulation (how much discomfort they experienced, how expressive they were, and how natural their reactions were) and the observers' attitude (how much empathy they felt for the demonstrator). Additionally, we asked the observers how well they could identify with the demonstrator. We asked the question in the pairs of friends only, as the pilot studies revealed problems with the question's understanding in the 'stranger' group (*n/a: no answer). Additionally, we asked the observers to rate the degree of unpleasantness attributed to the demonstrators (on a 1 - 5 scale). The last row shows comparisons between the 'friend' and 'stranger' groups, the Wilcoxon-Mann-Whitney test, p-values uncorrected. F - 'friend' group; S - 'stranger' group; IQR: interquartile range.

### 3. fMRI statistics

**Table S4**

*Activation Peaks in the Observational Learning Stage, US > no US Contrast*

| Cluster | Peak x | Peak y | Peak z | Peak value | Volume mm | Vox | Label |
|---|---|---|---|---|---|---|---|
| 1 | 50 | -66 | 6 | 14.98 | 89912 | 11239 | Right Lateral Occipital Cortex inferior division |
| | 44 | -76 | -8 | 12.92 | | | Right Lateral Occipital Cortex inferior division |
| | 42 | -44 | -16 | 12.21 | | | Right Temporal Occipital Fusiform Cortex |
| | 64 | -40 | 26 | 11.94 | | | Right Supramarginal Gyrus posterior division |
| | 52 | -28 | -6 | 11.74 | | | Right Middle Temporal Gyrus posterior division |
| | 50 | -40 | 10 | 11.51 | | | Right Supramarginal Gyrus posterior division |
| | 34 | 22 | -18 | 10.71 | | | Right Frontal Orbital Cortex |
| | 52 | 8 | -22 | 10.34 | | | Right Temporal Pole |
| | 22 | -88 | -8 | 10.07 | | | Right Occipital Fusiform Gyrus |
| | 46 | 26 | -4 | 9.82 | | | Right Frontal Orbital Cortex |
| | 42 | -2 | -14 | 9.71 | | | Right Planum Polare |
| | 52 | 24 | 22 | 9.14 | | | Right Inferior Frontal Gyrus pars opercularis |
| | 46 | 2 | 44 | 8.88 | | | Right Precentral Gyrus |
| | 12 | -68 | 38 | 8.81 | | | Right Precuneous Cortex |
| | 36 | -62 | -20 | 8.55 | | | Right Occipital Fusiform Gyrus |
| | 22 | -2 | -18 | 8.42 | | | Right Amygdala |
| | 14 | -98 | 6 | 8.17 | | | Right Occipital Pole |
| | 28 | -92 | 12 | 8.16 | | | Right Occipital Pole |
| | 38 | 6 | 4 | 7.50 | | | Right Insular Cortex |
| | 54 | 40 | -2 | 7.40 | | | Right Frontal Pole |
| | 64 | -32 | 40 | 7.35 | | | Right Supramarginal Gyrus anterior division |
| | 26 | -80 | 34 | 6.78 | | | Right Lateral Occipital Cortex superior division |
| 2 | -48 | -78 | 6 | 13.87 | 32568 | 4071 | Left Lateral Occipital Cortex inferior division |
| | -52 | -58 | 10 | 11.73 | | | Left Middle Temporal Gyrus temporooccipital part |
| | -62 | -46 | 22 | 11.32 | | | Left Supramarginal Gyrus posterior division |
| | -44 | -50 | -18 | 10.52 | | | Left Inferior Temporal Gyrus temporooccipital part |
| | -30 | -92 | -10 | 9.49 | | | Left Occipital Pole |
| | -24 | -66 | -28 | 6.64 | | | no label |
| | -28 | -94 | 14 | 6.22 | | | Left Occipital Pole |
| 3 | -32 | 20 | -14 | 10.65 | 15520 | 1940 | Left Frontal Orbital Cortex |
| | -30 | 28 | 2 | 8.87 | | | Left Insular Cortex |
| | -42 | 0 | -16 | 8.24 | | | Left Planum Polare |
| | -38 | -12 | -6 | 8.12 | | | Left Insular Cortex |
| | -20 | -4 | -14 | 7.58 | | | Left Amygdala |
| | -54 | 10 | 8 | 6.42 | | | Left Inferior Frontal Gyrus pars opercularis |
| 4 | 6 | 20 | 38 | 9.50 | 10368 | 1296 | Right Paracingulate Gyrus |
| | 4 | 20 | 56 | 9.07 | | | Right Superior Frontal Gyrus |

| | | | | | | | |
|---|---|---|---|---|---|---|---|
| | 6 | 40 | 10 | 8.55 | | | Right Cingulate Gyrus anterior division |
| | 10 | 4 | 72 | 6.61 | | | Right Superior Frontal Gyrus |
| 5 | -8 | -24 | -12 | 9.77 | 6816 | 852 | Brain-Stem |
| | 12 | -22 | -10 | 8.72 | | | no label |
| | 10 | -8 | 6 | 8.38 | | | Right Thalamus |
| | 10 | 8 | 4 | 7.85 | | | Right Caudate |
| | 6 | -4 | -10 | 7.13 | | | no label |
| 6 | -10 | -66 | 36 | 8.61 | 2336 | 292 | Left Precuneous Cortex |
| 7 | -2 | -24 | 28 | 8.69 | 2096 | 262 | Left Cingulate Gyrus posterior division |
| | 0 | -8 | 32 | 6.43 | | | Left Cingulate Gyrus anterior division |
| 8 | -16 | -94 | 26 | 7.12 | 1896 | 237 | Left Occipital Pole |
| | -26 | -78 | 24 | 5.95 | | | Left Lateral Occipital Cortex superior division |
| 9 | 34 | -52 | 50 | 6.78 | 1072 | 134 | Right Superior Parietal Lobule |
| 10 | -30 | -52 | 52 | 6.86 | 680 | 85 | Left Superior Parietal Lobule |
| 11 | 4 | 52 | 34 | 6.47 | 632 | 79 | Right Superior Frontal Gyrus |
| 12 | -10 | -76 | -44 | 6.58 | 440 | 55 | no label |
| 13 | -10 | -84 | 8 | 6.03 | 392 | 49 | Left Intracalcarine Cortex |
| 14 | -40 | -4 | 48 | 6.68 | 376 | 47 | Left Precentral Gyrus |
| 15 | -50 | -26 | -6 | 7.12 | 368 | 46 | Left Middle Temporal Gyrus posterior division |

*Note.* Table shows local maxima more than 16 mm apart. For brevity, clusters greater than 30 voxels were included. Last column gives the most probable label from the Harvard - Oxford atlas.

**Table S5**

*Activation Peaks in the Direct-expression Stage, CS+ > CS- Contrast*

| Cluster | Peak x | Peak y | Peak z | Peak value | Volume mm | Vox | Label |
|---|---|---|---|---|---|---|---|
| 1 | 30 | 30 | 0 | 8.30 | 5024 | 628 | Right Frontal Orbital Cortex |
| | 50 | 22 | 4 | 6.70 | | | Right Inferior Frontal Gyrus pars triangularis |
| 2 | -36 | 24 | -4 | 7.18 | 1400 | 175 | Left Frontal Orbital Cortex |
| 3 | -14 | -76 | -30 | 6.37 | 112 | 14 | no label |
| 4 | 10 | -12 | -12 | 6.25 | 64 | 8 | no label |
| 5 | 6 | 24 | 36 | 5.77 | 40 | 5 | Right Paracingulate Gyrus |
| 6 | 34 | 46 | 20 | 5.63 | 8 | 1 | Right Frontal Pole |
| 7 | 8 | 4 | 2 | 5.45 | 8 | 1 | Right Caudate |

*Note.* Table shows local maxima more than 16 mm apart. Last column gives the most probable label from the Harvard - Oxford atlas.

**Table S6**

*Activation Peaks for the Temporal Modulation of the CS+ Response in the Direct-Expression Stage*

| Cluster | Peak x | Peak y | Peak z | Peak value | Volume mm | Vox | Label |
|---|---|---|---|---|---|---|---|
| | | | | Both Groups | | | |
| 1 | -18 | -88 | -6 | 6.38 | 10480 | 1310 | Left Occipital Fusiform Gyrus |
| | -8 | -100 | 0 | 5.43 | | | Left Occipital Pole |
| | -12 | -80 | -32 | 4.49 | | | no label |
| | 10 | -86 | -10 | 4.47 | | | Right Lingual Gyrus |
| | 0 | -86 | 4 | 4.01 | | | Left Intracalcarine Cortex |
| | 24 | -76 | -14 | 3.65 | | | Right Occipital Fusiform Gyrus |
| | -26 | -98 | 20 | 3.58 | | | Left Occipital Pole |
| 2 | -4 | -54 | 62 | 5.42 | 5344 | 668 | Left Precuneous Cortex |
| | -12 | -76 | 44 | 4.84 | | | Left Lateral Occipital Cortex superior division |
| | 4 | -36 | 46 | 3.75 | | | Right Cingulate Gyrus posterior division |
| | -8 | -68 | 28 | 3.71 | | | Left Precuneous Cortex |
| 3 | 2 | 8 | 34 | 4.58 | 1704 | 213 | Right Cingulate Gyrus anterior division |
| | -2 | 26 | 24 | 3.66 | | | Left Cingulate Gyrus anterior division |
| 4 | 44 | 28 | 8 | 4.71 | 1184 | 148 | Right Inferior Frontal Gyrus pars triangularis |
| 5 | 6 | 46 | 32 | 5.67 | 848 | 106 | Right Paracingulate Gyrus |
| | | | | Friend > Stranger | | | |
| 1 | -58 | 14 | 10 | 5.83 | 2280 | 285 | Left Inferior Frontal Gyrus pars opercularis |
| 2 | -62 | -44 | 8 | 4.65 | 1328 | 166 | Left Supramarginal Gyrus posterior division |
| 3 | 16 | -58 | -2 | 4.41 | 1096 | 137 | Right Lingual Gyrus |
| 4 | -52 | 36 | 8 | 4.83 | 1072 | 134 | Left Inferior Frontal Gyrus pars triangularis |
| 5 | -24 | -6 | 12 | 4.55 | 768 | 96 | Left Putamen |

*Note.* Table shows local maxima more than 16 mm apart. Last column gives the most probable label from the Harvard - Oxford atlas.

**Table S7**

*Activation peaks for the psychophysiological interaction analysis based on the US > no US contrast*

| Cluster | Peak x | Peak y | Peak z | Peak value | Volume mm | Vox | Label |
|---|---|---|---|---|---|---|---|
| | | | | Anterior Insula | | | |
| 1 | 64 | -38 | 20 | 6.16 | 8520 | 1065 | Right Supramarginal Gyrus posterior division |
| | 54 | -48 | 4 | 5.00 | | | Right Middle Temporal Gyrus temporooccipital part |

| | | | | | | | |
|---|---|---|---|---|---|---|---|
| | 58 | -66 | 6 | 4.63 | | | Right Lateral Occipital Cortex inferior division |
| | 48 | -28 | 2 | 4.62 | | | Right Superior Temporal Gyrus posterior division |
| | 48 | -80 | 0 | 3.60 | | | Right Lateral Occipital Cortex inferior division |
| 2 | -52 | -56 | 12 | 5.26 | 6792 | 849 | Left Middle Temporal Gyrus temporooccipital part |
| | -42 | -70 | 12 | 4.80 | | | Left Lateral Occipital Cortex inferior division |
| | -68 | -46 | 8 | 4.01 | | | Left Middle Temporal Gyrus temporooccipital part |
| | -50 | -80 | -4 | 3.76 | | | Left Lateral Occipital Cortex inferior division |
| 3 | -12 | -74 | -14 | 4.88 | 3536 | 442 | Left Lingual Gyrus |
| | -30 | -74 | -22 | 4.64 | | | Left Occipital Fusiform Gyrus |
| | -40 | -64 | -14 | 3.66 | | | Left Occipital Fusiform Gyrus |
| 4 | 52 | 14 | 26 | 4.46 | 808 | 101 | Right Inferior Frontal Gyrus pars opercularis |
| 5 | 32 | -68 | -28 | 4.16 | 744 | 93 | no label |
| posterior STS | | | | | | | |
| 1 | 46 | -64 | 2 | 9.29 | 20656 | 2582 | Right Lateral Occipital Cortex inferior division |
| | 66 | -40 | 20 | 6.51 | | | Right Supramarginal Gyrus posterior division |
| | 38 | -80 | -14 | 5.01 | | | Right Lateral Occipital Cortex inferior division |
| | 46 | -40 | 10 | 4.64 | | | Right Supramarginal Gyrus posterior division |
| | 62 | -38 | 44 | 4.21 | | | Right Supramarginal Gyrus posterior division |
| | 32 | -94 | -4 | 3.51 | | | Right Occipital Pole |
| 2 | -46 | -70 | 8 | 8.07 | 12424 | 1553 | Left Lateral Occipital Cortex inferior division |
| | -58 | -48 | 12 | 5.20 | | | Left Supramarginal Gyrus posterior division |
| 3 | 38 | 32 | -2 | 5.68 | 4840 | 605 | Right Frontal Orbital Cortex |
| 4 | -58 | -40 | 36 | 4.70 | 3072 | 384 | Left Supramarginal Gyrus posterior division |
| | -66 | -32 | 24 | 4.63 | | | Left Supramarginal Gyrus anterior division |
| 5 | -38 | 22 | -8 | 5.01 | 2608 | 326 | Left Frontal Orbital Cortex |
| | -34 | 6 | -6 | 3.57 | | | Left Insular Cortex |
| 6 | 48 | 14 | 30 | 4.91 | 2240 | 280 | Right Inferior Frontal Gyrus pars opercularis |
| | 54 | 30 | 20 | 3.71 | | | Right Inferior Frontal Gyrus pars triangularis |
| 7 | 48 | -42 | -16 | 6.12 | 1992 | 249 | Right Inferior Temporal Gyrus temporooccipital part |
| 8 | 38 | -12 | -6 | 4.76 | 1768 | 221 | Right Insular Cortex |
| | 50 | -24 | -4 | 4.49 | | | Right Middle Temporal Gyrus posterior division |
| 9 | 12 | -2 | 12 | 4.93 | 1664 | 208 | Right Thalamus |
| | -2 | -4 | 4 | 3.94 | | | Left Thalamus |
| 10 | 54 | 20 | 8 | 4.96 | 1520 | 190 | Right Inferior Frontal Gyrus pars opercularis |
| 11 | 6 | 6 | 62 | 4.63 | 1344 | 168 | Right Juxtapositional Lobule Cortex (formerly Supplementary Motor Cortex) |
| 12 | 20 | 10 | -6 | 4.96 | 1120 | 140 | Right Putamen |
| 13 | -4 | -14 | -10 | 4.75 | 904 | 113 | no label |
| 14 | 6 | -24 | 0 | 4.49 | 816 | 102 | Right Thalamus |
| 15 | -40 | -62 | -28 | 4.65 | 744 | 93 | no label |
| posterior STS (small volume corrected within the amygdala) | | | | | | | |
| 1 | 24 | -4 | -16 | 3.82 | 104 | 13 | Right Amygdala |
| 2 | -22 | -4 | -14 | 3.56 | 24 | 3 | Left Amygdala |

| | | | | | | | |
|---|---|---|---|---|---|---|---|
| 3 | -20 | -6 | -18 | 3.42 | 8 | 1 | Left Amygdala |

*Note.* Table shows local maxima more than 16 mm apart. Last column gives the most probable label from the Harvard - Oxford atlas.
